# AutoSpect: An All-In-One Software Solution for Automated Processing of LA-ICP-TOF-MS datasets

**DOI:** 10.1101/2025.05.09.653091

**Authors:** Andrew M. Crawford, David Z. Zee, Qiaoling Jin, Aaron Sue, Niharika Sinha, Soo Hyun Ahn, Thomas V. O’Halloran, Keith W. MacRenaris

## Abstract

LA-ICP-TOF-MS provides rapid, high resolution elemental analysis of biological and non-biological samples. However, accurate real-time data analysis frequently requires the user to account for several instrumental and experimental variables that can change during data acquisition. AutoSpect is a novel software tool designed to automate the processing and fitting of LA-ICP-TOF-MS data, addressing key challenges such as time-dependent spectral drift, instrument sensitivity drift calibration inaccuracies, and peak deconvolution, enabling researchers to rapidly and accurately process complex datasets. The tool is optimized to be robustly applicable across scientific fields (e.g., geochemistry, biology, and materials science), providing a streamlined solution for end users seeking to maximize the potential of LA-ICP-TOF-MS for high-resolution elemental mapping and isotopic analysis.

**Significance to JAAS:** Analysis of fast transient signals using laser ablation inductively coupled plasma time-of-flight mass spectrometry (LA-ICP-TOF-MS) has become mainstream for elemental mapping. Advancements in LA-ICP-TOF-MS technology continue to accelerate the collective understanding of the role inorganic chemistry plays in dynamic processes. To ensure accurate quantitative results, the vast amount of complex spectral data generated requires elegant solutions to perform a variety of functions including data partitioning, peak fitting, drift correction, mass-to-charge calibration, peak profiling, and spectral fitting. AutoSpect is an all-in-one software solution that provides high level automation with a user-friendly graphical interface to perform complex data analyses for ICP-TOF-MS datasets.

## 1. Introduction

Time-of-flight mass spectrometry (TOF-MS) has fast become the technique of choice for analyzing fast transient signals following ionization via inductively coupled plasma (ICP) sources due to the dramatic improvement in high-speed detection electronics [1]. This has been especially true for single particle (SP), single cell (SC), and laser ablation (LA) ICP-TOF-MS applications ranging from nanoparticle and microplastic characterization to elemental mapping in geological and biological samples [2–4]. By combining high frequency LA with ICP-TOF-MS, rapid, high-resolution analyses at scales ranging from microns to millimeters are achievable. Additionally, with recent advancements in benchtop instrumentation, LA-ICP-TOF-MS is now capable of measuring all isotopes at frequencies from 10 Hz up to 1000 Hz helping to move elemental mapping into the realm of more routine, robust, quantitative analysis. These advancements are improving our fundamental understanding of how the distribution and concentration of elements play a role in dynamic processes and by combining elemental mapping with complementary imaging modalities such as MALDI-TOF-MS, NIR, Raman, histochemistry, immunofluorescence, and brightfield microscopy we can begin to unlock the complexity of a multitude of sample types [5–8].

The major advantage of ICP-TOF-MS is the ability to provide near simultaneous detection of all isotopes across the whole mass spectrum which makes it well-suited for the analysis of short transient events such as LA [9]. However, using TOF-MS for detecting these short transient events generated from single pulse-resolved laser ablation present challenges, including time-dependent spectral drift, instrument sensitivity drift, and the need for precise calibration and interference correction. Addressing these hurdles is crucial to ensuring accurate data interpretation, particularly for multi-element and isotopic analyses and quantitative elemental mapping. Although there are several software options for mass spectrometry image analysis there remain few commercially available spectral fitting tools for modern ICP-TOF-MS systems that handle spectral post-processing for ICP-TOF-MS datasets including LA [10]. Each of these tools have independent advantages but sequential application can lead to bottlenecks and inefficiencies in ICP-TOF-MS workflow. Most software [11–13] requires manual user input to identify optimal peaks for drift correction, mass axis calibration, and peak profile modeling, which can introduce subjectivity and user error. With larger specimens, a laser ablation imaging experiment can require several hours, exposing the spectra to greater possibility of drift due to external factors like temperature changes. While some commercially available software packages can accommodate peak drift up to a user-defined limit, but not automatically and can fail to accurately fit peaks that drift beyond that limit. Ideally, users would benefit from inspection of the accuracy of the drift correction process. Overcoming these challenges is particularly important when the goal is a quantitative comparison of dozens of analytes over a broad dynamic range in small regions of interest.

Herein, we describe the workflow of a new integrated and automated software solution called AutoSpect, that integrates multiple stages of data processing, beginning with raw data import, data sorting, peak identification, time dependent peak drift correction, mass-to-charge (*m*/*z*, or *μ*) calibration, peak profile creation, and spectral fitting to generate reliable outputs. Finally, we demonstrate AutoSpect use cases illustrating elemental map generation and data visualization as well as isotope accumulation.

## 2. AutoSpect Workflow

AutoSpect software provides a user-friendly graphical interface for analysis of complex ICP-TOF-MS datasets. This is accomplished through an automated process composed of the following six steps (**Figure 1**):

i. ***Processing and partitioning raw data:*** First, LA-ICP-TOF-MS data is imported in the raw hierarchical data format (HDF5), where each component is automatically defragmented to produce three distinct datasets (i.e. gas blanks, references, and samples). These three datasets are partitioned into hundreds of independently manageable data chunks with associated metadata (e.g., average integrated mass spectra, acquisition time, and m/z calibration coefficients). Data is then sorted and indexed into sample, gas blank, and references and partitioned to calculate averages.
ii. ***Measuring peak drift:*** Following processing of raw data, peaks are identified in the integrated spectrum and then tracked across spectral partitions to extract centroid positions. The centroids are modeled as smooth functions of acquisition time and used to characterize drift.
iii. ***Drift Correction:*** Using a reference set of centroids, partition-specific mass calibration parameters are optimized, modeled over time, and used to interpolate each partition onto the reference mass axis. Finally, a drift-corrected integrated spectrum is computed as a weighted average of the aligned partitions.
iv. ***Mass-to-charge (m/z) calibration:*** The global mass calibration parameters are recalibrated using a refined set of well-isolated peaks with known mass-to-charge (*m/z*) values. Each is fitted with Gaussian and Lorentzian models to extract centroids and widths. A smooth polynomial model is used to identify and iteratively remove outliers, leaving a final set of peaks for recalibrating the global parameters. The parameters are then refined iteratively until convergence, and the updated calibration parameters are propagated to update drift correction results.
v. ***Peak profiling:*** Empirical peak profiles are generated based on isolated peaks that are automatically identified by AutoSpect. These profiles are used to create a look up table (LUT) of all possible peaks contained in the range of the spectra. This ensures accurate representation of spectral features corresponding to the main peak centers. AutoSpect then accounts for instrumental notching effects and models peak tailing.
vi. ***Spectral fitting:*** The generated peak profiles are applied to fit spectra, resolving overlapping peaks and deconvoluting interferences. Nonlinear optimization algorithms refine the fitting process, ensuring high precision in determining elemental and isotopic abundances. Fitted spectra are saved and a new HDF5 file is generated that can be used in a variety of downstream image processing programs such as iolite.

**Figure 1.**
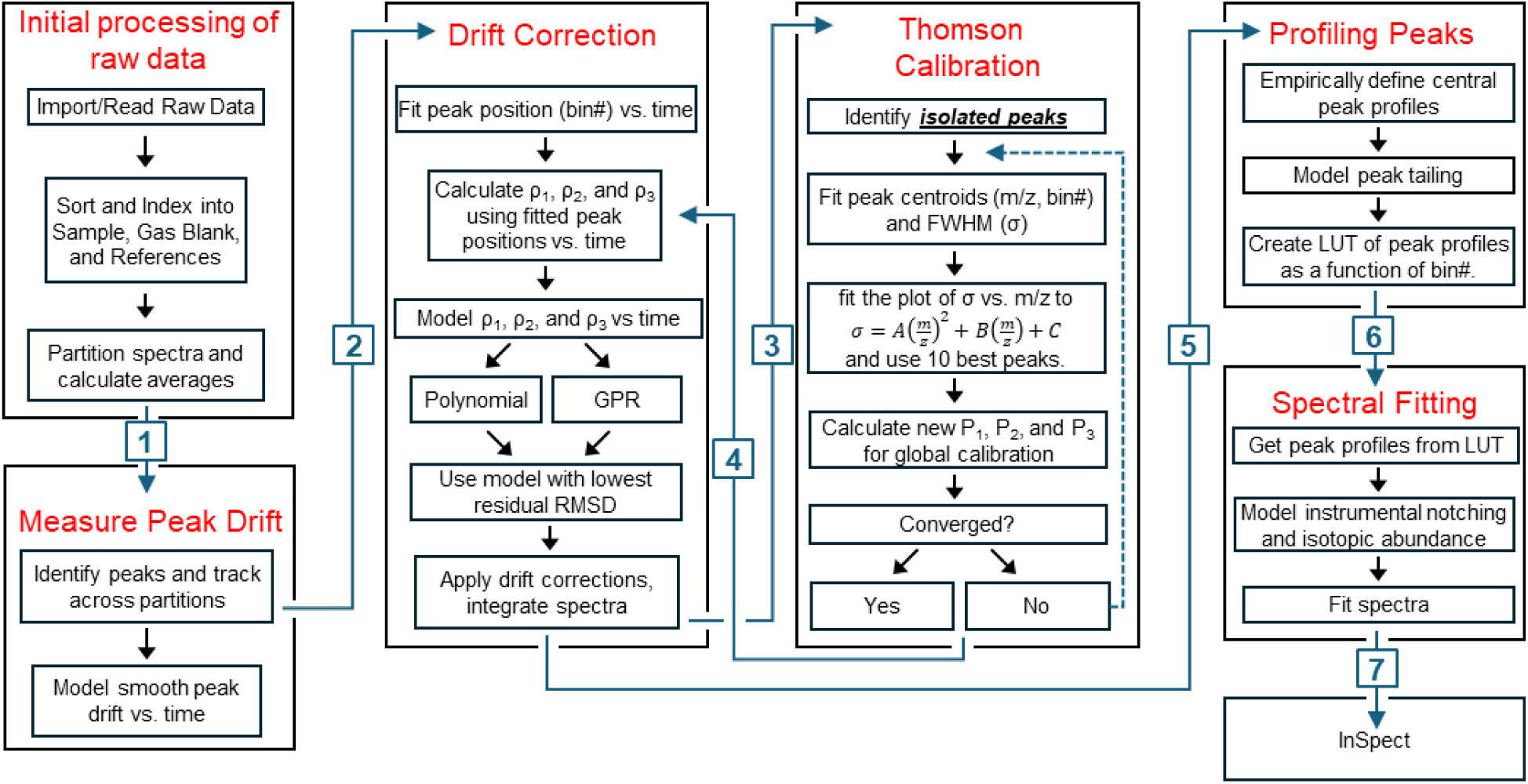
AutoSpect Workflow. AutoSpect was designed as an all-in-one software solution for ICP-TOF-MS datasets that performs many processes automatically. For contextualization, the processes have been broken into six sections. Following initial processing of the raw data, AutoSpect measures the peak drift across all measured spectra. Once peak drift has been measured, the drift correction is calculated and applied. Using the drift corrected integrated spectrum the mass-to-charge axis is recalibrated, and then the drift corrections are updated to the optimized axis. Using the drift corrected and *m/z* optimized spectra, AutoSpect empirically determines the peak profiles for all possible *m/z* contained in the data. Finally, AutoSpect performs fitting using linear least squares to deconvolute all peaks and produce elemental mapping data.

## 3. Materials and Methods for AutoSpect use cases

Datasets to test the key functionality of AutoSpect such as peak fitting and branching were obtained from fresh frozen murine kidney and epididymis as well as sea urchin egg samples. Mice organs were dissected from Balb/c and C57BL/6 mice and were embedded in optimal cutting temperature (OCT) media in a plastic mold and flash frozen using chilled isopentane over liquid nitrogen as described elsewhere [14]. Briefly, samples were sectioned at 10 µm using a conventional cryostat (Leica Biosystems) onto SuperFrost charged slides (Thermofisher Scientific) and stored in −20°C prior to ablation. Sea urchin egg from *Arbacia punctulata* (Marine Biology Laboratory, Woods Hole, MA, USA) were isolated following standard procedures [15], deposited onto Kapton thin film (SPEX SamplePrep) and allowed to air dry at ambient temperature for 8 hours prior to ablation.

Experimental setup begins with outlining the regions of interest on the sample (i.e. kidney, epididymis), Additionally, the instrument is set to measure at least 5 seconds of gas blanks. This ensures appropriate collection of signals that can be used for background subtraction.

All samples were ablated using the Elemental Scientific Lasers BioImage 266 nm laser ablation system (Elemental Scientific Lasers, Bozeman, MT, USA) equipped with an ultra-fast low dispersion TwoVol3 ablation chamber and a dual concentration injector and connected to icpTOF S2 mass spectrometer (TOFWERK AG, Thune, Switzerland). Elemental content was analyzed real time according to *m/z* ratio. Tuning was performed using NIST SRM612 glass certified reference material (National Institute for Standards and Technology, Gaithersburg, MD, USA). Torch alignment, lens voltages and nebulizer gas flow were optimized using ^140^Ce and ^55^Mn while maintaining low oxide formation based on the ^232^Th^16^O+/^232^Th+ ratio (<0.5). Murine samples were ablated in a raster pattern with a laser repetition rate of 100 Hz using a 10 µm circular spot. Sea urchin samples were ablated in a raster pattern with a laser repetition rate of 100 Hz, a 2 µm circular spot, and zero overlap. Scanning data was recorded using TofPilot 1.3.4.0 (TOFWERK AG) and saved in the HDF5 format. Laser ablation images were visualized using Iolite [16,17].

## 4. AutoSpect Software

In a time-of-flight mass spectrometer (TOF-MS), ions are accelerated through an electric potential and then travel through a field-free drift region before striking the detector. The time that an ion takes to reach the detector is determined by its mass-to-charge ratio (m/z). Since each ion undergoes the same accelerative force, the more massive an ion is the slower the acceleration it undergoes. This separates the ions in space based on their mass, such that the corresponding digital spectral time bin, *b_i_*, for a particular μ_*i*_ is given by Equation 1 and the corresponding μ_*i*_ for a given time bin, *b_i_*, is given by Equation 2.

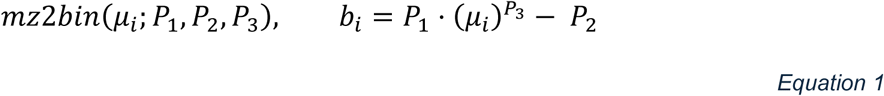

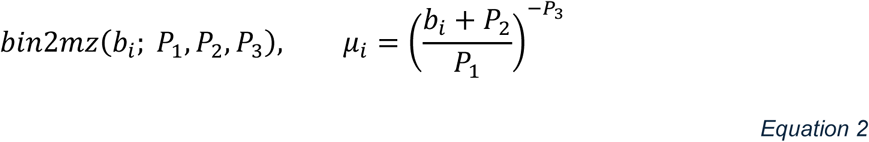

where P_1_, P_2_, and P_3_ are fixed coefficients determined during instrument calibration prior to data collection, and routinely recalibrated after a scan is collected.

### 4.1 Partitioning and processing raw data

Laser ablation based elemental mapping is acquired using line scans. As such, a single image dataset is represented by as many line scans as there are rasters in the image. Thus, a single image corresponds to the aggregate of thousands of individual line scans. Furthermore, as an LA-ICP-TOF-MS instrument collects an image, it alternates between segmented scans of the sample, reference standards, and gas blanks. As the instrument alternates between segmented scans of the sample, reference standards, and gas blanks, the resulting spectra are fragmented and dispersed across numerous acquisitions. To facilitate structured data handling, spectra from all measurements— sample, gas blank, and references—are sorted accordingly and grouped into smaller partitions, each containing approximately 1600 spectra.

For each partition, the average acquisition time and spectrum are computed and stored. The spectral data is structured into a partition matrix, ***S*** ∈ ℝ^***N***×***Q***^, where N represents the number of spectral bins, and Q denotes the number of partitions. Each column ***S***(: , *q*) in the matrix corresponds to a partition containing the mean spectrum for that subset:

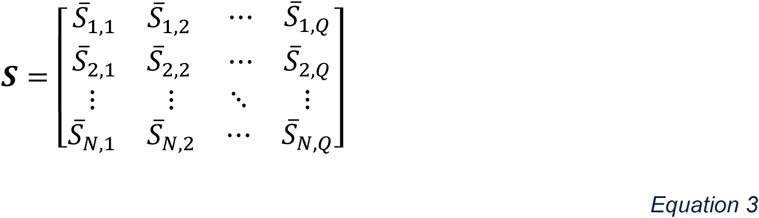

where each element *S̅*_*n*,*q*_ represents the average intensity at spectral bin *n* within partition *q*. The partition-wise averaged spectrum is then computed (**Equation S1**). The columns of ***S*** are then sorted in ascending order of their mean acquisition times, ***t*** ∈ ℝ^***Q***^, with the *q*^th^ entry of ***t*** corresponding to the *q*^th^ column of ***S***. Lastly, the integrated spectrum, ***s*** ∈ ℝ^***N***^, is obtained by averaging over all individual raw pixel spectra.

### 4.2 Peak Identification, Tracking Peaks, and Measuring Peak Drift

TOF instruments can be sensitive to fluctuations in physical parameters such as temperature. This can lead to time-dependent peak centroid drift of the ions of interest over the course of sample analysis. As such, the peak centroids’ positions at the initiation of a scan may not reflect their corresponding positions throughout the scan. Therefore, this time-dependent peak drift must be accounted for before any processing/fitting of the resultant spectra to produce the most accurate isotope maps. The major drawback for peak fitting is the subjective nature of removing peaks prior to fitting and the time constraints of full spectrum peak fitting. AutoSpect solves these issues through development of an automated process to identify peaks and measure peak drift.

#### Peak Identification

To track and measure peak drift, AutoSpect identifies peaks in the integrated spectrum and then tracks across spectral partitions to extract centroid positions. Using the integrated spectrum, peaks are identified as local maxima where the intensity decreases monotonically to the nine nearest neighboring bins on both sides. The set criteria are satisfied to ensure that the center bin is a true local maximum with symmetrical monotonic decay in intensity over a 19-bin window (see **Equations S2** and **S3**). Using the integrated spectrum, peaks are identified as local maxima where the intensity decreases monotonically to the nine nearest neighboring bins on both sides.

#### Tracking peaks across spectral partitions

Each spectral partition, ***S***(: , *q*), is analyzed as follows. The intensity values for each peak region in ***S***(: , *q*) are extracted and assembled into a matrix **Λ**_*q*_ ∈ ℝ^**19**×***J***^, where each column, *j*, contains the raw intensity values 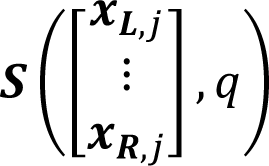 such that

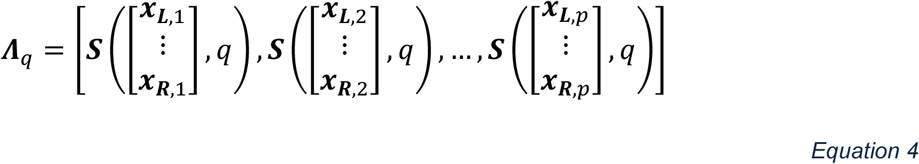

Here, each column in **Λ**_*q*_ represents the 19-bin peak region, centered at ***x***_*j*_. Similarly, the corresponding bins for the intensity values of the peak regions in ***S***(: , *q*) are extracted and assembled into a matrix **Λ**_*x*_, *i* ∈ ℝ^**19×J**^, such that

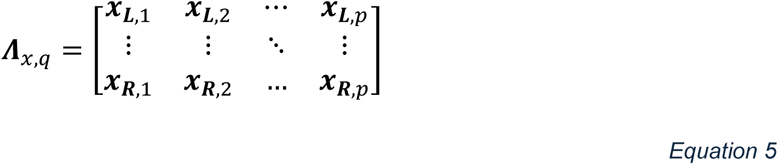

Each peak region **Λ**_*q*_(: , *j*) is first corrected by subtracting a linear baseline computed using a first-order polynomial fit. The baseline is subtracted from the raw peak region, yielding a corrected peak region. This baseline-corrected region is then fitted to a Lorentzian function. The optimal parameters are determined by minimizing the sum of squared residuals (See **Section 3** in Supporting Information).

From each spectral partition, *q*, and peak region, *j*, the fitted centroid, *x*_0_, is stored to the *q*-th and *j*-th entry of matrix ***x***^_**0**_ ∈ ℝ^***J***×***Q***^, where ***J*** is the number of peaks and ***Q*** is the number of spectral partitions (see **Figure 2A**, gray lines).

**Figure 2.**
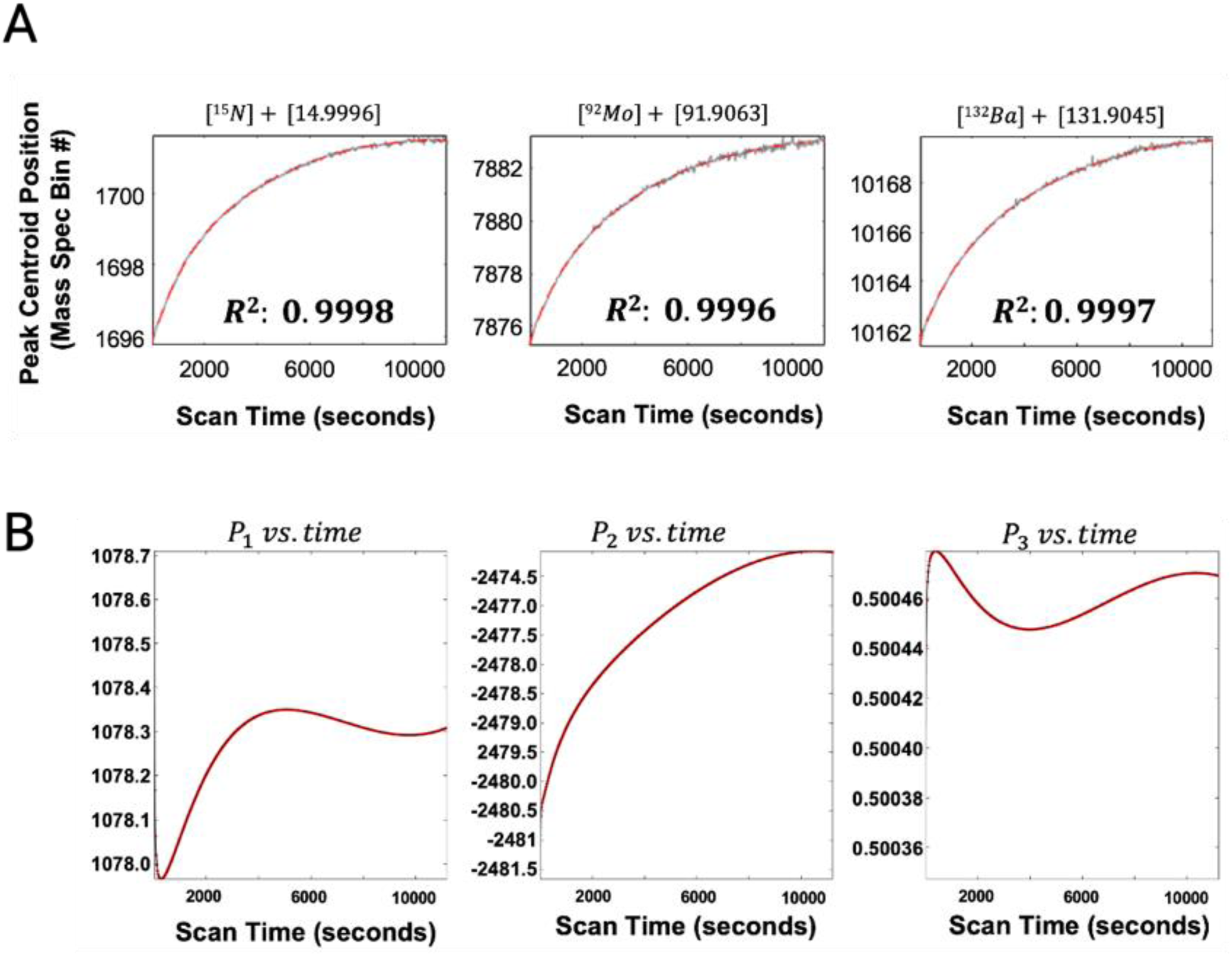
AutoSpect conducts robust automatic drift correction with error less than 1 ppt. **(A)** Plots of measured peak centroids for ^15^N, ^92^Mo, and ^132^Ba as a function of time from each partitioned average spectrum showing 3 of the 20+ accepted peaks to be used in calculating drift correction. **(B)** Following peak tracking and identification, the fitted peak centroids are used to calculate the coefficients for the Thomson calibrations necessary for drift correction. The red dots = fitted/measured centroid; black line = fitted theoretical values for P_1_, P_2_, and P_3_ for the whole scan. They are superimposed to one another.

#### Solving a smooth function of peak position vs. time

Where ***x***^_**0**_ is the array of fitted centroids from our Lorentzian modeling to the peaks across the spectral partitions, we seek to solve for ***x***^_***f***_, the smooth temporal trend of the centroids across the partitions. To do this, each row of the peak centroid matrix, ***x***^_**0**_, is independently fitted to a smooth function of the mean acquisition time, ***t***, (**Figure 2A**, red fitted line, Equation S10). Using the R^2^ calculated from the fit to each row (peak centroid as a function of time), only the rows corresponding to the top 20 R^2^ values are retained and used in the following section to solve spectral drift as a function of acquisition time. This ensures that the most temporally coherent and well-modeled peaks are used for spectral drift correction.

### 4.3 Modeling Peak Drift and Applying Drift Correction

#### Selecting the Basis Set or Reference Spectrum

Since the peak centroids shift as a function of time, each spectral partition has its own unique set of coefficients used for Equation 1 and Equation 2. To account for the drift, AutoSpect models the coefficients for each partition as a function of acquisition time. This creates three new variables, **ρ**_**1**_, **ρ**_**2**_, and **ρ**_**3**_ ∈ ℝ^***Q***×**1**^, each representing a vector of length Q corresponding to the spectral partition specific entries for ***P***_**1**_, ***P***_**2**_, and ***P***_**3**_, respectively (**Equation S11**).

AutoSpect solves for the coefficients **ρ**_**1**_, **ρ**_**2**_, and **ρ**_**3**_, by first assigning a basis set for comparison. This basis set provides a reference set of peak centroids for comparison with the time-dependent peak centroids derived above. This set of basis peaks is determined by assigning one of the partitions as a reference. To determine the reference partition, total peak region counts per spectral partition are computed and the partition with the fourth-highest total counts is chosen as the reference.

#### Calculating Partition Specific Mass Calibration Parameters

To refine the mass calibration parameters, AutoSpect sets about solving the partition specific parameters, stored in the vectors ***ρ*_1_**, ***ρ*_2_**, and ***ρ*_3_**, used for interconverting between bin# and *m/z (*μ) (**Equation 1** and **Equation 2**). Using the *j*^*tℎ*^ column index of the ***x***^_*f*_, the peak centroids are converted from bin# to μ (**Equation 2**) and become the basis set against which the rest will be compared.

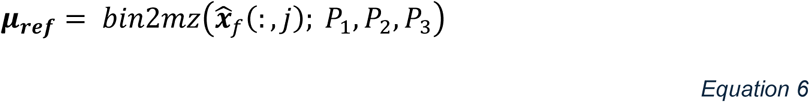

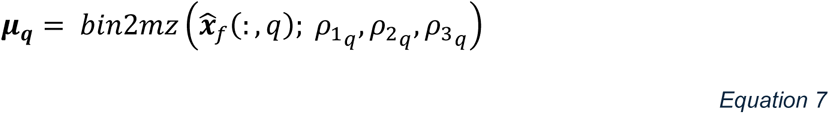

Using the theoretical centroids calculated above, ***x***^_*f*_, the partition specific calibration parameters (**ρ**_**1**_, **ρ**_**2**_, and **ρ**_**3**_) are solved by minimizing the residuals between expected/basis (μ_***ref***_, Equation 6) and fitted (μ_*q*_, Equation 7) values.

Following optimization of the **ρ**_**1**_, **ρ**_**2**_, and **ρ**_**3**_calibration parameters, each parameter is fitted as a function of the mean acquisition time, ***t***, using polynomial fitting, Gaussian Process Regression fitting, or a simple cubic spline interpolation. The resulting root-mean-squared-deviations from each approach for all three parameters are compared and the approach with the corresponding lowest value is used going forward (**Figure 2B**).

Once the spectra of ***S*** are drift corrected, the integrated spectrum is recalculated (**Figure 3** bottom row, **Figure 6a**) as the weighted mean of the drift corrected spectral partitions.

**Figure 3.**
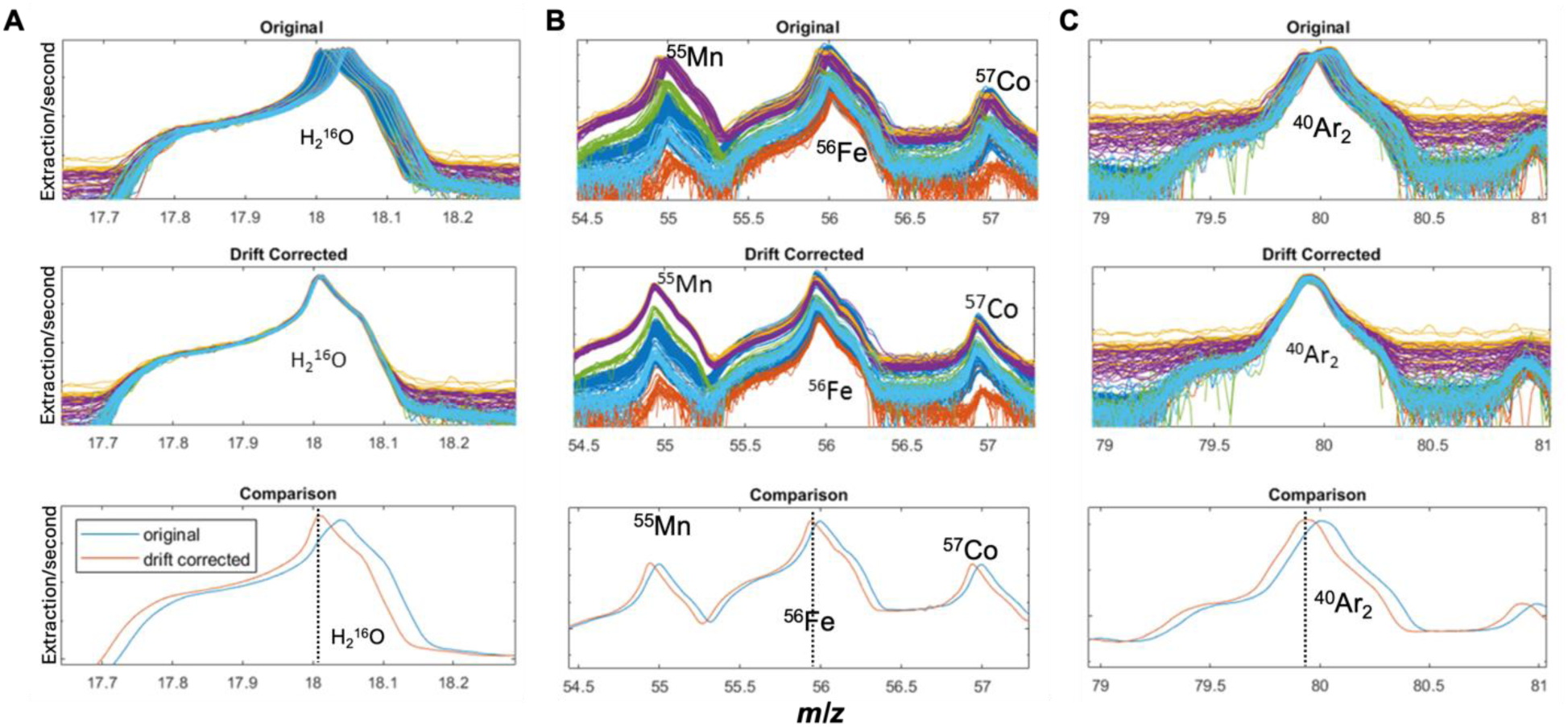
Time dependent instrument drift correction. AutoSpect automatically corrects for drift, which is applied to the whole dataset. Here we show drift correction for five masses (**A**) H_2_^16^O, (**B**) ^55^Mn, ^56^Fe, ^57^Co, and (**C**) ^40^Ar_2_. The top panel shows the original uncorrected spectra, the middle panel shows drift corrected spectra, and the bottom panel shows a comparison of the average of both the original (blue) and drift corrected (red) spectra. Both plots show significant changes to the apparent peak position and peak morphology post AutoSpect.

#### Drift correction of spectral partitions

Each spectral partition ***S***(: , *q*) is corrected for time-dependent drift using the optimized partition-specific calibration parameters **ρ**_**1**_(*q*), **ρ**_**2**_(*q*), and **ρ**_**3**_(*q*) for each spectral partition *q* = 1, . . . , *Q*. Each partition is first converted to its corresponding mass axis, μ_*q*_, using Equation 7, with subsequent interpolation from μ_*q*_onto the reference axis μ_***ref***_using a shape-preserving cubic Hermite operator (Equation 8).

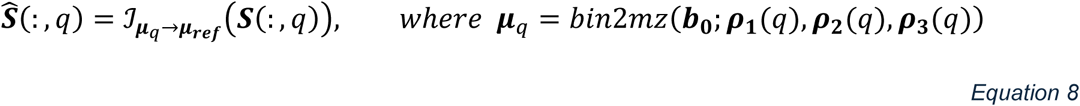

Here:

- ***0***_**0**_ is the uniformly spaced vector 1, …, *N*, corresponding to bins of μ_***ref***_,
- μ_*q*_ is the mass axis computed from ***x***_0_ using Equation 2 for partition *q*,
- ℐ_μ*q*→μ***ref***_ is the interpolation operator, implemented using a piecewise cubic Hermite interpolation polynomial, that maps the spectrum from the local axis μ_*q*_ to the global reference axis μ_***ref***_,
- **S**^(: , *q*) is the drift-corrected spectrum matrix for partition *q*.

An overlay of the various spectral partitions following drift correction can be seen in **Figures 3A and 3B**, which show drift correction of low, mid, and high range *m/z* regions. This can be seen by the top and middle rows. Where the top rows show noticeable drift in the centroid positions, the middle row shows that the drift corrected spectra are perfectly aligned.

#### Recomputing the Integrated Spectrum

Peak drift results in blurred features from the integrated spectrum. To correct for this blurring, the integrated spectrum must be recalculated following drift correction. Since each partition is derived from a known number of raw spectra, the drift-corrected integrated spectrum is then computed as a weighted average of the columns of **S**^, with weights proportional to the number of contributing spectra (**Section 8**, Supporting Information).

The deblurring of spectra following drift correction can be seen in **Figure 3** by looking at the bottom row of plots and comparing the original integrated spectrum with the integrated spectrum from drift correction. Although the peaks are narrower for all three regions, the impact, at least in this example, is most noticeable for *m/z* of 18, where it is easily seen that the peak is much more clearly defined.

### 4.4 Optimizing Mass Spectrum Calibration

The global mass calibration parameters are recalibrated using a refined set of well-isolated peaks with known *m/z* values. Each is fitted with Gaussian and Lorentzian models to extract centroids and widths. A smooth polynomial model is used to identify and iteratively remove outliers, leaving a final set of peaks for recalibrating the global parameters. The parameters are then refined iteratively until convergence, and the updated calibration parameters are propagated to update drift correction results.

#### Identifying isolated peaks for global mass spectrum calibration

Following drift correction, the global variables *P*_1_, *P*_2_, and *P*_3_ are recalibrated using actual mass peaks, i.e., both a peak exists and the corresponding value of its μ is known (rather than assumed) with very high confidence to be a specific ion (e.g. ^55^Mn). This is different from drift correction where only the presence of the peak and its apparent μ according to reference spectrum and the suboptimal mass calibration coefficients *P*_1_, *P*_2_, and *P*_3_ set by the instrument were used.

This begins by using the preset integration windows for the 315 ions in the look-up table (LUT) available from TofWare. All peaks that have integration windows that do not overlap with any other window are considered isolated peak regions. For the icpTOF S2 instrument, of the 315 available peak regions in the LUT, 118 peaks are considered isolated. Then in a process similar to peak identification above (Section 4.2), each peak region of the drift corrected weighted integral spectrum, ***s***^, is checked to verify two things.

1. The highest point (peak) is contained in the window but not at the boundaries of the window.
2. The spectral intensities are monotonically decreasing to the left and right (similar to **Equations S2** and **S3**).

### Fitting Isolated Peak Regions to obtain the Centroids and Peak Widths

Following initial identification, the isolated peak regions are analyzed using Gaussian and Lorentzian fitting approaches. For each peak region, the raw intensity values, *y*_*i*_, and corresponding indices, *x*_*i*_, are extracted along with the corresponding *m/z* values for the corresponding ion, μ_*i*_^*LUT*^, from the LUT. The region defined by the index corresponding to the point of maximum intensity and a window of ±4 is fitted to a single Gaussian and the R^2^ calculated. Then only peak regions with R^2^>0.7 are carried forward for further analyses.

Next, the fitted centroid and width from the Gaussian fitting are used as initial values for fitting the same region to a Lorentzian function (**Equation S17**, Supporting Information). From this, the centroids (***x***_***fitted***_) and peak widths (**σ**_***fitted***_) are saved. To avoid spurious fits, the ***x***_***fitted***_ is verified to be contained within the windowed region. If it is not, then the peak region and its associated ***x***_***fitted***_ and **σ**_***fitted***_ are discarded. Using the R^2^ from the Lorentzian fitting, all fitted peak regions with R^2^>0.75 are carried forward. Using Equation 2, the ***x***_***fitted***_ is converted to μ_***fitted***_. Then ***x***_***fitted***_, μ_***fitted***_, and **σ**_***fitted***_ are stored into column vectors for further refinement (Equation S18, Supporting Information).

#### Solving for Peak Broadening as a Function of *m/z*

Using μ_***fitted***_ along with **σ**_***fitted***_, the list of isolated peaks is further refined to determine the optimal peaks to be used for recalibrating the global coefficients, ***P***_**1**_, ***P***_**2**_, and ***P***_**3**_. The peak widths (σ) follow a smooth trend as a function of the mass-to-charge ratio (μ) [13]. Since individual peaks may not always be fully isolated, i.e. some may be broadened due to overlap with neighboring peaks, a filtering process is applied to iteratively remove peaks with broadened widths. If a peak overlaps with another, the fitting process may incorrectly attribute a broader width to what is a combination of multiple peaks. By removing outliers with excessive width, we improve confidence that the remaining peaks are truly isolated and that their widths accurately reflect instrumental resolution and physical broadening effects.

To model the expected smooth dependence of **σ**_***fitted***_ on the corresponding fitted centroid positions μ_***fitted***_, a second-order polynomial model is constructed using non-negative least squares regression. In addition, the expected peak width for each centroid is then estimated using a fitted polynomial model then the most extreme outlier is removed, and the polynomial fit is recomputed with the remaining peaks. This process is repeated until only 10 peaks remain (**Figure 4A**, Supporting Information **Section 10**).

**Figure 4.**
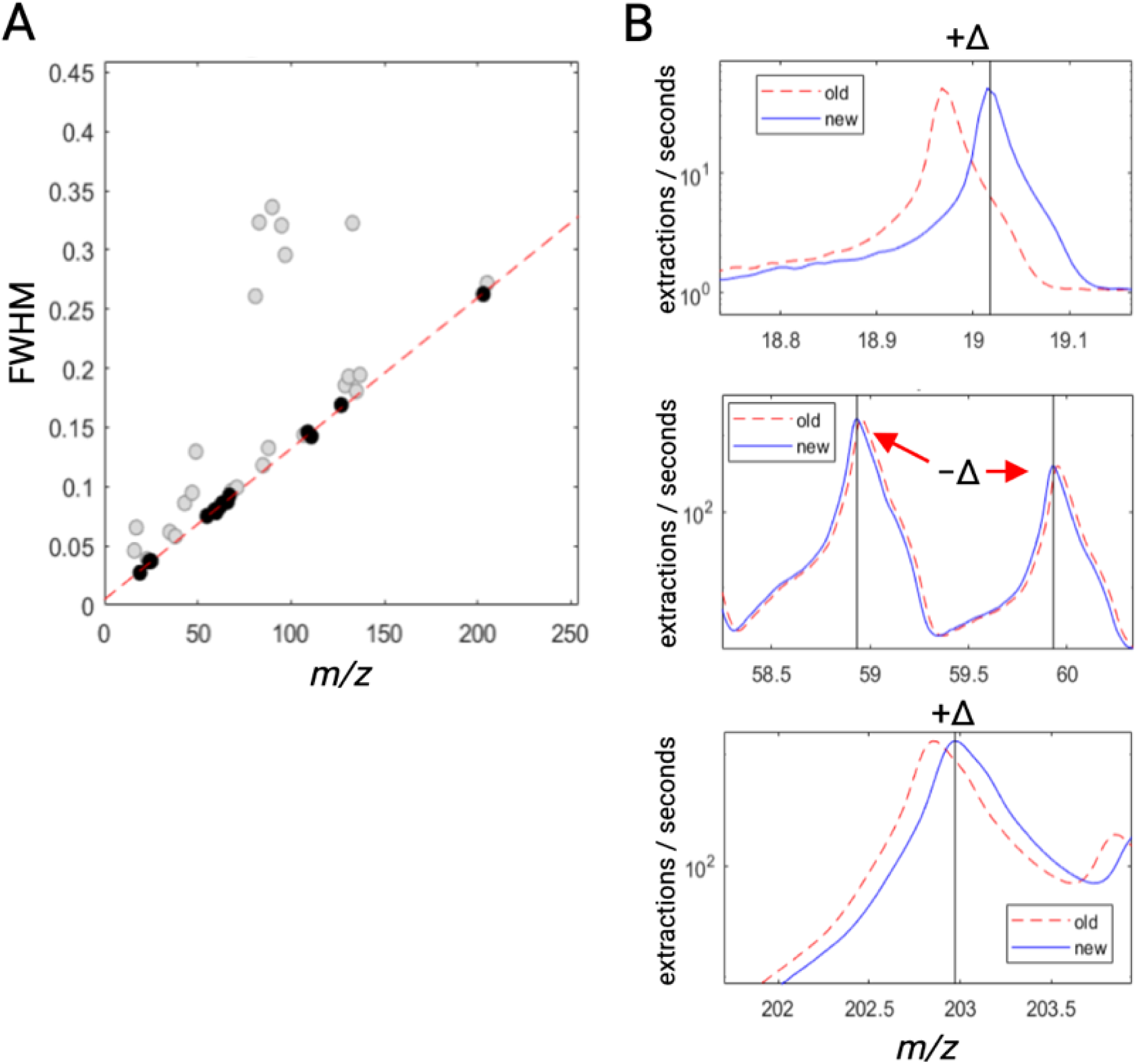
Final mass calibration. **(A)** Fitted peak widths are modeled as a function of *m*/*z* because isolated ions (in black) should exhibit narrower peak widths than isobars (in grey). This relationship is used to recalibrate the spectra post collection. Using nonlinear optimization with nonnegativity constraints, the plot of **σ** vs. **μ** is fit to the equation **σ=**a**μ^**2+b**μ**+c. The peaks used for fitting are black dots with light grey representing peaks that were rejected during analysis. **(B)** Four different peaks from three different spectral regions (*m*/*z* 18.8 – 19.1, 58.5 – 60, 202 – 203.5) comparing the average spectrum pre (red, dashed line) and post drift correction (blue, solid line).

From these 10 peaks, the associated ***x***_***fitted***_ and μ_***fitted***_ are used to optimize the coefficients ***P***_**1**_, ***P***_**2**_, and ***P***_**3**_. The optimization problem is solved using nonlinear least squares (**Equation S20**, Supporting Information). The associated **σ**_***expected***_ from the ten peaks are used to calculate the expected peak centroid width for all spectral bins for use in **sections 4.5** and **4.60**.

#### Iterative Refinement of Mass-to-Charge Axis Recalibration

Post calibration of drift correction parameters, the *m/z* axis for the detected spectra needs to be recalibrated. The recalibration can be conducted from the integrated spectrum, but the peaks are blurred because the integrated spectrum is the average of drifted peaks. Mass-to-charge axis recalibration is performed after drift correction using the highly resolved peaks. Refinement of mass calibration is an iterative process. Each iteration begins with the mass axis (*M*_*i*_) determined by the initial values for the coefficients *P*_1*i*_, *P*_2*i*_, and *P*_3*i*_, and ends with optimized values for the coefficients (*P*_1*i*+1_, *P*_2*i*+1_, and *P*_3*i*+1_) and *M*_*i*+1_. At the end of each iteration, the unique values from *M*_*i*_ and *M*_*i*+1_are taken combined and the combined bin positions using the previous and current parameters are computed. Then, recalibration is considered converged when the population becomes self-consistent (**Section 11**, Supporting Information). **Figure 4B** shows how various regions of the spectra can shift, and that recalibration of the *m/z* axis is not always just a simple translation. As can be seen in **Figure 4B**, around both *m/z* of 19 and 203, both regions required a corrective shift to higher *m/z*, while around *m/z* of 60 required a corrective shift to lower *m/z*.

#### Post recalibration optimization of drift correction

The global calibration coefficients **P_1_**, **P_2_**, and **P_3_** are used in drift correction as the basis set for calculating the *m/z* axis into which the peaks from all the spectral partitions are aligned. Post-drift-correction optimization of these parameters means the spectral partition coefficients for **ρ**_**1**_, **ρ**_**2**_, and **ρ**_**3**_(**Equation S11**, Supporting Information) must also be updated. Therefore, the optimization procedures performed in **section 0** are repeated using the new set of **P_1_**, **P_2_**, and **P_3_**.

### 4.5 Peak Profile Calculations

#### Dilation of Isolated Peak Regions

The 10 isolated peak regions, identified in **section 4.4 0** are adaptively dilated. The left boundary (***x***_***L***_) of the integration window is initially set at the starting index assigned by the LUT for each identified peak. To adjust this boundary, AutoSpect iteratively examines the mean intensity in a forward window (***x***_***L***_ to ***x***_***L***+**3**_) and a backward window (***x***_***L***−**3**_ to ***x***_***L***_). If the mean intensity in the forward window exceeds that of the backward window, the boundary is shifted leftward by one unit, i.e., ***x***_***L***_ → ***x***_***L***−**1**_. This process continues until the intensity trend reverses or the boundary approaches the lower limit of the spectrum, at which point further shifting is halted, and the final ***x***_***L***_ is assigned. Similarly, the right boundary (***x***_***R***_) is initially set at by the LUT and adjusted dynamically. If the mean intensity in the forward window (***x***_***R***_ to ***x***_***R***+**3**_) is lower than that in the backward window (***x***_***R***−**3**_ to ***x***_***R***_), the boundary is shifted rightward by one unit, i.e., ***x***_***R***_ → ***x***_***R***+**1**_. This adjustment continues until the trend reverses, or the boundary reaches the upper spectrum limit, at which point further shifting ceases, and ***x***_***R***_ is set (**Section 12**, Supporting Information).

#### Constructing an Empirical Peak Center Profile Look Up Table

The isolated peak regions spanning the full *m/z* range are each normalized to an intensity range spanning from 0 to 1. The peaks are then transformed from *m/z* space into standard deviation space, centered at the fitted centroid μ₀ and scaled by the fitted Gaussian peak width σ:

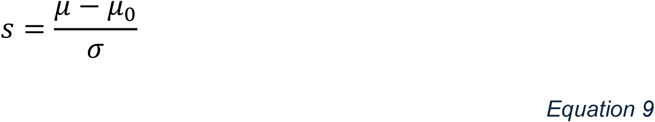

Each transformed peak is then interpolated onto a fixed standard deviation grid, *s* ∈ [−20, 20], with k uniformly spaced points. The peak profiles corresponding to a centroid positioned anywhere on the mass-axis are interpolated across all detector bins using first-order polynomial regression at each s-coordinate, forming a continuous 2D surface of peak shape profiles. This creates the empirical look-up tables (LUT) used below:

- LUT_*y*_ ∈ ℝ^*k*×*N*^: empirical peak profile intensities in standard deviation space, with each column corresponding to the measured profile at each of the N integer value digital spectral bins.
- LUT_*x*_ ∈ ℝ^{*k*^ ^×^ ^*N*}^: empirical peak profile spectral bin indices in standard deviation space, with each column corresponding to the bin coordinate of the corresponding intensity value in LUT_*y*_.

This LUT facilitates accurate, calibration-driven construction of empirical peak shapes during fitting.

#### Empirical Peak Center Profile via LUT Interpolation

Given a desired centroid *m/z* denoted as μ₀ ∈ ℝ^m^, for each of m peaks, the corresponding empirical peak shape is retrieved via interpolation from the precomputed LUTs. This process involves several variables, an interpolation operator, and a series of transformation steps (**Section 13**, Supporting Information).

#### Interpolation Procedure

The empirically determined peak profiles are determined from our LUTs as follows. First, *m/z* is converted to fractional bin index, *b̅*, using the calibration parameters using Equation 1 and global calibration coefficients (*P*_1_, *P*_2_, and *P*_3_). Since *b̅* will only rarely be a whole integer (i.e., an exact match to a column of LUT_*x*_ and LUT_*x*_), the two nearest corresponding integer bins are blended to generate the intermediate profile corresponding to *b̅* . Then, using *x* and *y* , the peak shape *y*_i_ , defined over non-integer spectral bin coordinates *x*_*i*_, in standard deviation space, is mapped onto the uniform detector bin grid via interpolation and then normalized to unit area. This yields *G* ∈ ℝ^*N*×*m*^, the empirical core peak shapes for each centroid μ_0,*i*_.

#### Asymmetric Tailing Functions

Visual inspection of the peaks in mass spectra show long tails that extend to low and high *m/z* by tens of *m/z* units (**Figure 5B-D**). **Figure 6** shows the result of accounting for these tails and the impact they can have on very distant *m/z* peaks. To account for these low and high *m/z* extended tails, each peak’s core shape is extended with asymmetric leading and trailing tails similar to those modeled in x-ray fluorescence microscopy [18–20].

**Figure 5.**
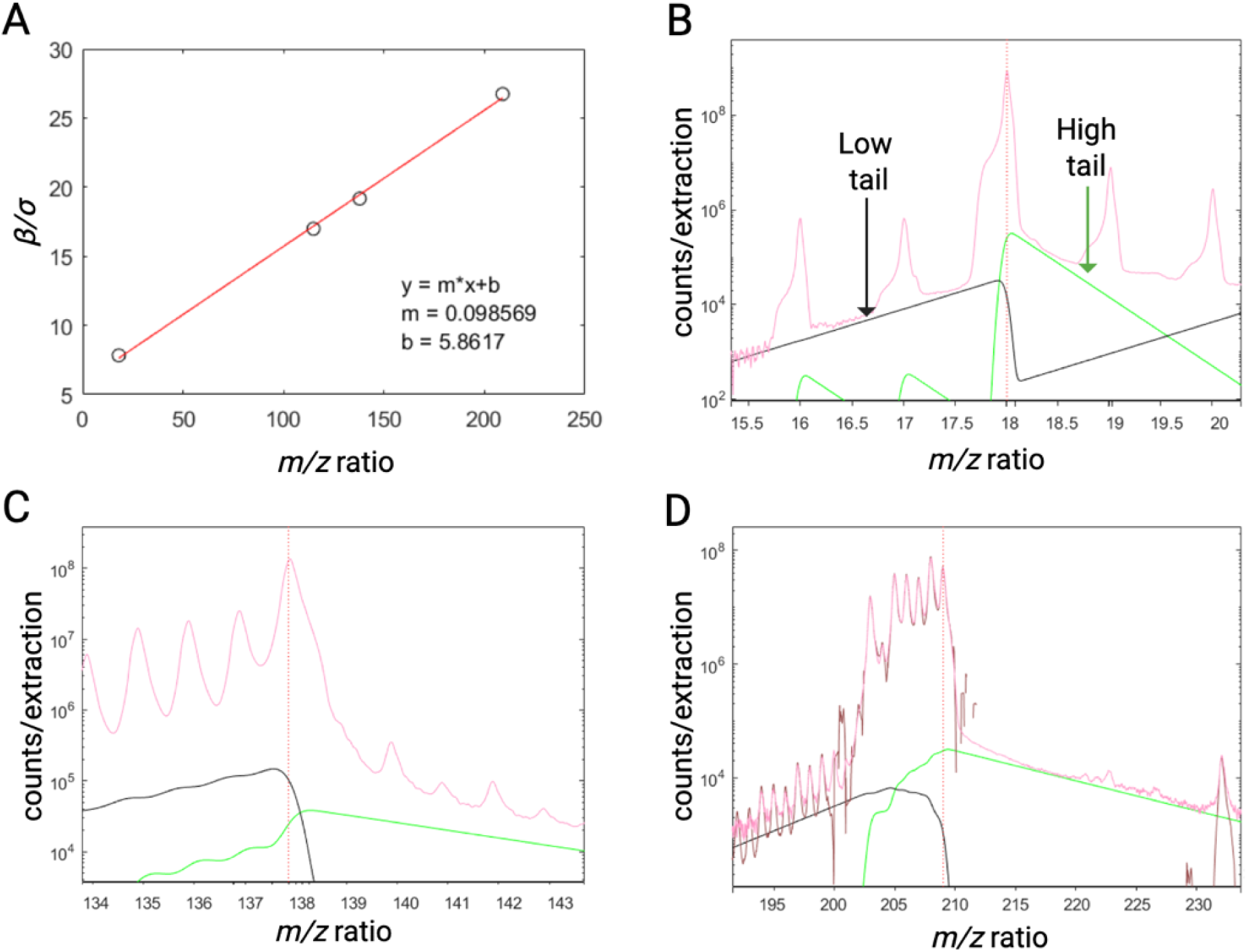
Peak tailing. **(A)** High and low peak tails are fitted to an error function and the width (β) is constrained to a ratio of the centroid peak width (σ) fitted by a Lorentzian. The ratio *β/σ* is constrained to a smooth function of *m/z* (here we show a 1^st^ order polynomial). **(B-D)** Shows the calculated high and low peak tails for ^1^H_2_^16^O^+^ (B), ^138^Ba^+^ (C), and ^208^Pb^+^. Of note, this is the same equation used in x-ray fluorescence to model low energy tailing of the fluorescence peaks caused by inefficient electron capture in the detector, or residual energy. For LA, there is a very steep low *m/z* but a long trailing high *m/z* tail. The high *m/z* is where expected detector or focusing inefficiencies manifest since higher masses arrive later.

**Figure 6.**
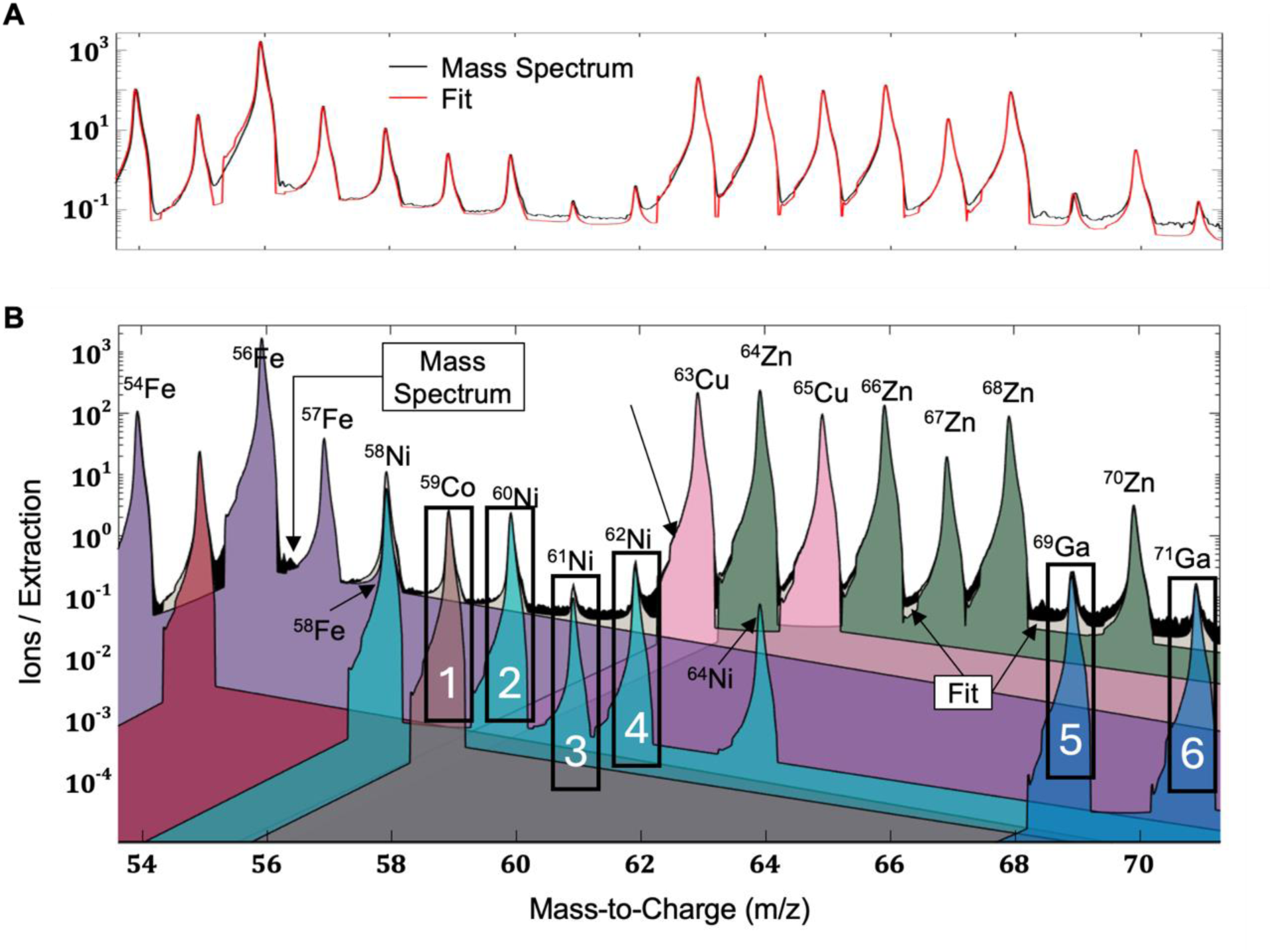
The importance of tailing in accurately fitting peaks across the spectra. **(A)** The integrated spectrum fit of a murine brain tissue section. **(B)** An area plot of the overlapping mass spectrum isotopes from *m/z* 54 to 71 displayed on a log intensity scale. The spectra were corrected by gas blank measurement and the residual fitted to the parameterized peak profiles automatically determined in AutoSpect. The fit to the empirically determined profiles with isotope ratio branching are shown for Mn, Fe, Co, Ni, Cu, Zn, and Ga. The peak centroids are not fully empirically determined which is why they drop to the low and high *m/z* tails.

Let:

- μ ∈ ℝ^*N*×1^: m/z evaluation points
- σ ∈ ℝ^1×*m*^: Peak widths
- ***β*** ∈ ℝ^1×*m*^: tail widths

The tails are defined as:

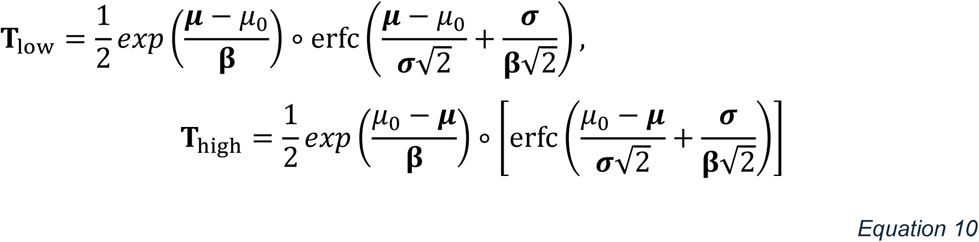

Each tail matrix is normalized column-wise (**Equation S34**, Supplemental Information).

#### Tail Widths Constrained by Peak Width

Tailing widths, ***β***, follow a smooth dependence on peak resolution across the mass range. As such, tail widths are not free parameters; rather, they are scaled by the peak widths according to a first-order polynomial in centroid *m/z* (**Section 15**, Supporting Information). This relationship between ***β*** and **σ** and the strong adherence to a first order relationship can be seen in **Figure 5A** which shows the ratio of ***β***/**σ** as a function of *m/z*.

#### Mass-to-Charge Dependent Tail Amplitude Scaling

The amplitude of the low- and high-*m/z* tails are on the order of 0.0001 of the intensity of the parent peak and vary smoothly across the spectrum (**Section 16**, Supporting Information). This scaling function is applied independently to the low- and high-energy tails to produce the vectors ***r*_low_** and ***r*_high_**, which are then used below during peak shape assembly.

#### Peak Shape Combination and Normalization

After empirical peak shapes and asymmetric tails have been defined, they are combined into a final asymmetric model, G_T_, for each elemental signal. This step ensures that each modeled peak incorporates core instrument resolution and realistic tailing. After combining the core peak with the asymmetric tails, each resulting peak shape *G*_T_(: , *i*) is normalized such that its total area integrates to one (**Equation S34**, Supporting Information). The resulting matrix G_T_ ∈ ℝ^*N*×*m*^ contains the asymmetric, resolution-constrained, and normalized peak shapes for each elemental signal prior to application of instrumental notching and isotopic branching.

#### Correction for Instrumental Notching (Tofwerk-Specific)

AutoSpect includes an optional correction mechanism specific to Tofwerk TOF-MS instruments, which employ inverse quadrupole filtering, referred to as notching, to reduce the transmission of extremely abundant background ions, such as those arising from argon, nitrogen, water vapor, and other atmospheric components, which would otherwise cause serious detector saturation. This filtering alters the transmission efficiency at selected *m/z* values, artificially suppressing the observed intensities of certain isotopes and thereby distorting their natural abundance ratios.

To correct for this effect, AutoSpect uses the stored notching parameters from the raw data file to automatically optimize and account for these changes. These values are used to compute a set of scaling factors, referred to as FLOAT, which are applied to the branching ratios of affected isotopes.

#### Isotopic Ratio Branching Application

Since the isotopes of given elements have very well documented natural abundances, the ratios of the various isotopes for a given element can be used to combine the equations for individual isotopes of a given species into a single equation accounting for the split probabilities of each isotope. If a sample contains isotopic enrichment, that isotope can be disconnected from branching and fitted individually.

To model isotopic structure, each elemental peak in **G**_*T*_ is condensed into one or more isotopic peaks using the notch-corrected branching ratio matrix **B̃**_iso_.

Let:

- **G**_T_ ∈ ℝ^***N***×***m***^: asymmetric, tailed, normalized elemental peak profiles,
- **B̃**_iso_ ∈ ℝ^***m***×γ^: corrected branching ratio matrix,
- **G**_br_ ∈ ℝ^***N***×γ^: final modeled isotope-resolved asymmetric, tailed peak profiles.

Then, the isotopic peak shapes are computed as:

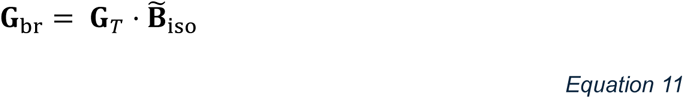

Each column of **B̃**_iso_ contains fractional contributions from isotopes of a given element. These are derived from known natural abundances and adjusted to account for notching. Each column is then normalized to unit area (**Equation S34**, Supplemental Information). This step allows each species to be represented by a single equation, incorporating all isotopes and correcting for instrument-induced artifacts. It improves both interpretability and robustness in spectral deconvolution, particularly for peaks that are partially resolved or overlapping (**Figure 6**).

### 4.6 Spectral Fitting and Peak Deconvolution

In the final steps of AutoSpect, using the drift calibration and the branched equations defined in the previous sections, the spectra for the samples, gas blanks, and reference standards are fitted to deconvolute peak overlap. Typically, with LA data, the individual pixel spectra are sparsely populated where most of the spectral bins contain zeros. This is especially the case with biological tissues. Because of this, interpolating the pixel spectra into a common grid for drift correction can result in significant errors. To avoid such potential errors, the composite model matrix *G*_*br*_ is interpolated to the local drift-corrected mass axis for each partition. Then, prior to spectral fitting, background correction is applied using either measured gas blanks or an automated SNIP-based background estimation algorithm. Once the spectra and background corrected and the model matrix is aligned with the measured data, the corrected spectra are deconvolved using standard linear least squares:

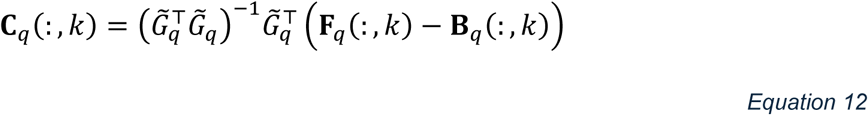

This yields ***C_q_***, a matrix of coefficients for all components (columns of *G̃*_*q*_) across all spectra in partition *q*. Since each column of *G̃*_*q*_ is normalized to unit area, the resulting coefficients in ***C***_*q*_ represent the total extracted counts for each modeled peak component, and no additional normalization or post-processing is required (**Section 19**, Supporting Information).

Once fitted, AutoSpect copies the original data file and then updates fitted peak data. Since AutoSpect outputs the data in the native TofWerk format, the fitted data can be seamlessly incorporated to any of the existing data reduction pipelines currently in the field using the TofWerk hdf5 structure.

## 5. AutoSpect Use Case Examples

### Single Cell and Tissue Section Imaging

AutoSpect functionality was tested on freshly isolated sea urchin eggs from *Arbacia punctulata*, which have an average diameter of 70 um). This dataset required ∼around 4.5 hours per sample and exhibited noticeable peak drift (**Figure 8**). Data sets on 10 µm thick epididymis and kidney tissue sections that showed low and high localized abundance of molybdenum, respectively, an essential enzymatic cofactor.

AutoSpect was compared to currently existing software for spectral analysis, Tofware, to benchmark performance. A laser ablation dataset of sea urchin eggs was analyzed using both Autospect and Tofware to compare their workflows, speed, and performance. This dataset was chosen due to its large file size, with 987 raster lines of 1305 spots, each 2 µm in size, giving a field of 2610 × 1974 µm, and a file size of 9.08 GB. The Tofware workflow totaled 53 min and 24 s, comprising peak shape optimization and peak fitting across 315 masses ranging from ^6^Li+ to ^238^UO+ (see **Table 2**). The analogous peak fitting by Autospect was accomplished in 47 min 7 s. The Autospect workflow took 39 min 17 s to complete drift correction, and an additional 1 min 44 s to perform peak profiling, totaling 1 h 28 min 8 s for all operations. The resulting fitted laser ablation images and the original, uncorrected data are shown in **Figure 8**. Notably, this dataset has fluctuating P and Zn signals due to environmental factors and/or peak tailing over the course of the analysis. AutoSpect eliminates these fluctuations and gives a stable response over the course of the experiment.

**Table 1.** The contribution of the various tails to the separate isotope peaks are calculated for six regions identified in **Figure 6**. Although the area of any tail negligibly contributes to the quantitative estimates of its parent isotope, these tails have very far-reaching impacts across the data. This can be seen by comparing the relative amplitudes of the various tails with the total peak amplitude for the various isotopes.

| REGION | PEAKS |  | Tails |  |  |  |  |  |
| --- | --- | --- | --- | --- | --- | --- | --- | --- |
|  |  |  | Intensity |  |  | % of Peak |  |  |
|  | Isotope | intensity | Fe | Cu | Zn | Fe | Cu | Zn |
| 0 | 56Fe | 1526 | 0.2408 | - | - | 0.02% | - | - |
| 1 | 59Co | 2.526 | 0.0998 | - | - | 3.95% | - | - |
| 2 | 60Ni | 2.291 | 0.0725 | - | - | 3.16% | - | - |
| 3 | 61Ni | 0.1542 | 0.0520 | 0.0027 | - | 33.74% | 1.76% | - |
| 4 | 62Ni | 0.3078 | 0.0380 | 0.0092 | - | 12.33% | 2.99% | - |
| 5 | 69Ga | 0.2553 | 0.0039 | 0.0106 | 0.0283 | 1.52% | 4.15% | 11.08% |
| 6 | 71Ga | 0.1671 | 0.0020 | 0.0063 | 0.0175 | 1.22% | 3.76% | 10.45% |

**Table 2.** Comparing the data processing tasks and the times needed by AutoSpect and Tofware to complete those tasks. Spectral data processing with AutoSpect and Tofware were performed on the same machine.

| <u>Data processing task</u> | <u>Time taken by AutoSpect</u><br><u>(hh:mm:ss)</u> | <u>Time taken by Tofware</u><br><u>(hh:mm:ss)</u> |
| --- | --- | --- |
| Dataset 1: sea urchin eggs (experimental scan time = 4:37:15) |  |  |
| peak fitting<sup>a</sup> | 0:47:07 | 0:53:24 |
| drift correction | 0:39:17 | —<sup>b</sup> |
| peak profiling | 0:01:44 | —<sup>b</sup> |
| total | 1:28:08 | 0:53:24 |
| Dataset 2: tissue slide 2 (experimental scan time = 1:10:02) |  |  |
| peak fitting<sup>a</sup> | 0:05:39 | 0:07:18 |
| drift correction | 0:17:33 | —<sup>b</sup> |
| peak profiling | 0:00:17 | —<sup>b</sup> |
| total | 0:23:29 | 0:07:18 |
| Dataset 3: tissue slide 3 (experimental scan time = 1:10:04) |  |  |
| peak fitting<sup>a</sup> | 0:04:45 | 0:09:12 |
| drift correction | 0:19:05 | —<sup>b</sup> |
| peak profiling | 0:00:14 | —<sup>b</sup> |
| total | 0:24:04 | 0:09:12 |
| Dataset 4: tissue slide 5 (experimental scan time = 1:20:54) |  |  |
| peak fitting<sup>a</sup> | 0:09:09 | 0:11:06 |
| drift correction | 0:24:12 | —<sup>b</sup> |
| peak profiling | 0:00:20 | —<sup>b</sup> |
| total | 0:33:41 | 0:11:06 |

<sup>a</sup>Includes data loading and writing tasks.
<sup>b</sup>function not available

Another demonstration for AutoSpect involves showing the effectiveness of isotope ratio branching in revealing spatial localization of low isotopic abundance elements such as molybdenum in biological tissues (**Figure 7**). Molybdenum acts as an essential metal cofactor to enzymes that participate in redox reactions due to its ability to shuttle between three oxidation states [21]. Therefore, detection of Molybdenum in biological samples most likely indicate the presence of these enzymes. While Molybdenum was detectable using Iolite software, the isotope ratio branching capability of AutoSpect improved signal to noise, allowing for accurate assessment of localization within the sample.

**Figure 7.**
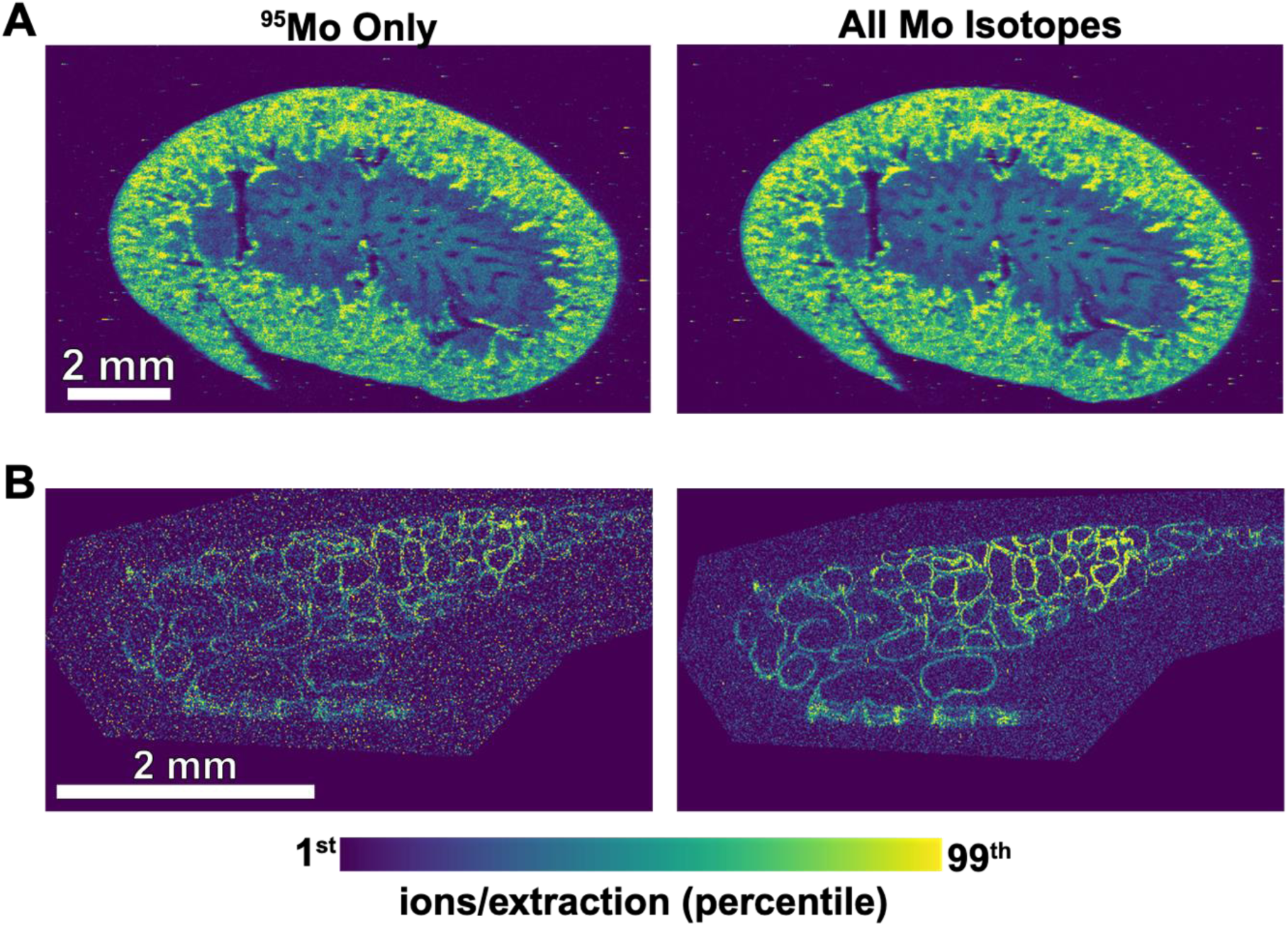
Isotope ratio branching improves signal to noise for low abundance elements in biological tissue samples. **(A)** LA-ICP-TOF-MS Molybdenum map of a 20 µm thick mouse kidney section. Kidney has naturally high concentrations of Mo and does not benefit as much from isotope ratio branching. However, **(B)** LA-ICP-TOF-MS of a 20 µm thick mouse epididymis, a male reproduction organ, which has naturally low concentrations of Molybdenum shows how mapping using a single isotope (^95^Mo) has lower signal to noise than if isotope ratio branching branching (combining other isotopes of Mo with variable abundance), improves signal to noise allowing accurate localization of the element within the tissue.

**Figure 8.**
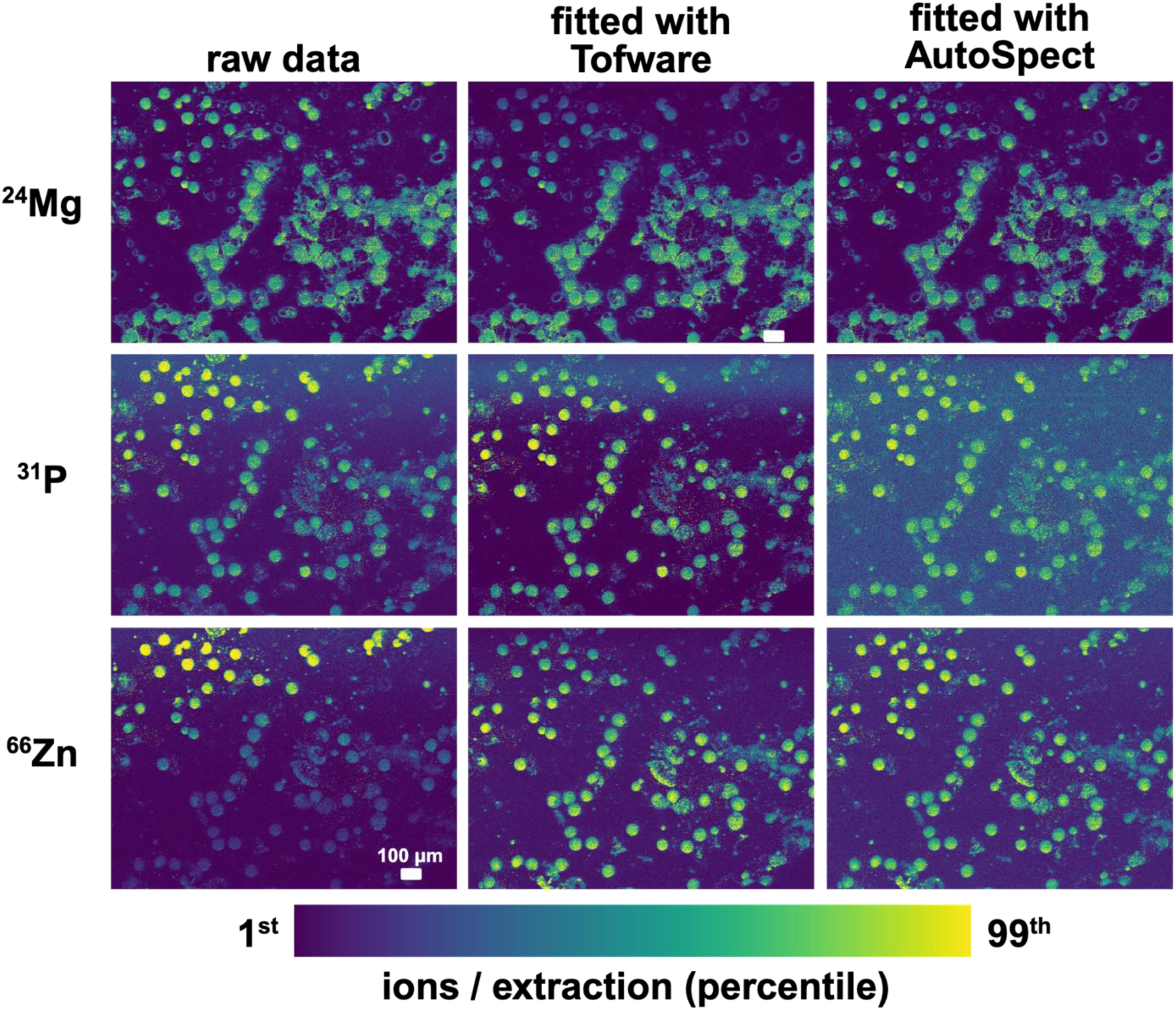
Comparison of peak fitting and drift correction by AutoSpect versus Tofware. Sea urchin eggs were deposited onto Kapton thin film and analyzed by LA-ICP-TOF-MS. The ^24^Mg, ^31^P, and ^66^Zn data are visualized from the 1^st^ to 99^th^ percentile. The resulting, unprocessed data showed localization of ^24^Mg, ^31^P, and ^66^Zn in the eggs (first column). Notably, the ^31^P and ^66^Zn signals fluctuate during 4.5 hours of acquisition experiment—we tentatively attribute these fluctuations to interferences with the tails of ^16^O_2_ (m/z = 32) and ^40^Ar (m/z = 40) gas peaks, respectively, at the beginning of the scan. When fitted with Tofware, the eggs have better contrast from background in the ^31^P and ^66^Zn channels, but all three mass channels show fluctuation (second column). Fitting with AutoSpect can eliminate these artifacts and give a stable response over the course of the experiment (third column).

## 6. Discussion

AutoSpect provides a transformative approach to LA-ICP-TOF-MS analysis, addressing longstanding challenges in data processing. By automating drift correction and peak profile determination, profiling low and high peak tailing, and fitting spectra, it improves both the accuracy and efficiency of analysis. AutoSpect’s flexible design allows for adaptation to other fields requiring high-resolution elemental and isotope analyses. Future updates will incorporate additional functionality, such as advanced multiply charged species handling and enhanced matrix-matching algorithms.

The approach employed in AutoSpect, and many of the underlying algorithms, were built upon ideas developed for x-ray fluorescence microscopy (XFM) [22–25]. In particular, the implementation of branching ratios, introduction of the low and high *m/z* tailing, use of directly subtracting the gas blank, and the partitioning and chunking of spectra were adapted from XFM.

### Recalibration of the Mass Axis

When identifying isolated peaks to calculate the smooth function of peak broadening as a function of *m/z*, the field largely supports the use of a first order polynomial. However, a second order polynomial was used rather than a first order (**Section 10**, Supporting Information) as it was found that allowing for subtle curvature adhering to a square component yielded peaks that aligned with their tabulated *m/z* values across the dynamic range of individual atoms to a much better degree than a slope component alone (**Figure 4A**).

### Automatic generation of peak profiles

Empirical peak profiles are generated based on isolated peaks that are automatically identified by the software and used to create a LUT of all possible peaks contained in the range of the spectra. This ensures accurate representation of spectral features corresponding to the main peak centers. AutoSpect then accounts for instrumental notching effects and models peak tailing. The incorporation of peak tailing and accounting for these tails is quantitatively important, especially concerning the high *m/z* tailing which persists for tens of *m/z*. This explains the difficulty of fitting an isotope neighboring another isotope with an intensity that is greater than 4 orders of magnitude larger.

### Peak tailing

Peak tailing is a new feature that has not been implemented into peak shape modeling of mass spectra. The functional forms have been adapted from XFM for modeling inefficient electron capture in detectors. For example, the contribution from the peak tail for ^56^Fe to be 0.02% the amplitude of the main centroid contributing negligibly to the quantitative estimate of ^56^Fe. However, because LA exists across a dynamic range of 8 to 10 orders of magnitude, these tails can have significant non-negligible impacts on neighboring peaks near and far in unpredictable manners that are sample dependent. For example, comparing the amplitudes of the raw peak and tails present for ^61^Ni (region 3), 35.5% of the peak amplitude corresponds to the combined contributions from the Cu and Fe tails (**Figure 6**). Similarly, 15.32% of the peak amplitude for ^62^Ni (region 4) corresponds to these two tails. For ^69^Ga (region 5) and ^71^Ga (region 6), 16.76% and 15.43% of the peak amplitudes correspond to the combined contributions of the Fe, Cu, and Zn tails. By comparison, the 3.95% and 3.16% contributions to ^59^Co (region 1) and ^60^Ni (region 2), are less significant.

### Spectral Fitting

AutoSpect applies the generated peak profiles to fit spectra, resolving overlapping peaks and deconvoluting interferences. Nonlinear optimization algorithms refine the fitting process, ensuring high precision in determining elemental and isotopic abundances. Fitted spectra are saved for further analysis and reporting.

## 7. Conclusion

AutoSpect represents a significant advancement in the field of LA-ICP-TOF-MS, providing researchers with a robust, automated tool for spectral fitting and data reduction. Case use studies indicate that AutoSpec supports:

1. Enhanced spectral deconvolution, resolving interferences with high precision.
2. Accurate drift corrections, ensuring consistent results across extended analytical sessions.
3. Efficient handling of complex datasets, reducing analysis times significantly compared to manual methods.

By streamlining workflows and addressing critical analytical challenges, AutoSpect empowers users to fully leverage the capabilities of this powerful analytical technique, paving the way for new discoveries in fields from geology to biology, from medicine to material sciences.

## 8. Author contributions

AMC conceived, designed, developed, wrote, and validated all software, including AutoSpect, InSpect, and affiliated subroutines, and performed testbed analyses with input from KWM, DZZ, and TVO. AMC also contributed to the interpretation of results and preparation of the methods description. AMC, DZZ, SHA, KWM, TVO wrote the manuscript. DZZ performed software comparisons, isolated samples, collected and analyzed results for the sea urchin data sets. Mouse kidneys were harvested by QJ. KWM, DZZ, SHA, NS, and AS sectioned murine tissues and collected data.

## 10. Data availability

The data supporting the findings of this study are available within the article and its ESI.† Data and software are available upon request and can be found at https://qemap.ehi.msu.edu/resources.

## Acknowledgements

Metal analysis and quantitative elemental mapping was performed at the Michigan State University Quantitative Bio Element Analysis and Mapping (QBEAM) Center generously supported by the MSU Office for Research and Innovation, the National Institute of General Medical Sciences of the National Institutes of Health under award number P41 GM135018 and by the Office of The Director, National Institutes of Health under award number S10OD026786. D.Z.Z. thanks the National Institute of General Medical Sciences of the National Institutes of Health for a Ruth L. Kirschstein NRSA postdoctoral fellowship (F32GM139401). SHA acknowledges support from National Institute of Child Health and Disease R01 HD100832 for supporting acquisition of mouse epididymal tissue, and TVO acknowledges R01 GM038784 for supporting sea urchin samples. The content of this publication is solely the responsibility of the authors and does not necessarily represent the official views of the National Institutes of Health.

