## supplemental for "AutoSpect: An All-In-One Software Solution for Automated Processing of LA-ICP-TOF-MS datasets"

### Supporting Information

**Section 1**: where each element $\bar{S}_{n,q}$ represents the average intensity at spectral bin $n$ within partition $q$. The partition-wise averaged spectrum is computed as:

$$\bar{S}_{n,q} = \frac{1}{K_q} \sum_{m=1}^{K_q} S_{n,m}^{(q)}$$

*Equation S1*

where $K_q$ is the number of individual spectra in partition $q$, and $S_{n,m}^{(q)}$ represents the raw spectral intensity for bin $n$ in $m^{th}$ spectrum of partition $q$.

The columns of $\boldsymbol{S}$ are then sorted in ascending order of their mean acquisition times, $\boldsymbol{t}$, of size $Q \times 1$ with the $q^{\text{th}}$ entry of $\boldsymbol{t}$ corresponding to the $q^{\text{th}}$ column of $\boldsymbol{S}$.

Lastly, the integrated spectrum, **s**, a column vector of length *N*, is obtained by averaging over all individual raw spectra from all scans (i.e., sample, gas blanks, and reference standards).

**Section 2 (Peak Identification)**: Using the integrated spectrum, $\boldsymbol{s}$, peaks are identified as local maxima where the intensity decreases monotonically to the nine nearest neighboring bins on both sides. Specifically, for each candidate peak, centered at bin $\boldsymbol{x}_j$, the following conditions must be satisfied:

$$\boldsymbol{s}(x_n) < \boldsymbol{s}(x_{n+1}) \; \forall x_n \, \epsilon \left[\boldsymbol{x}_j - 9, \boldsymbol{x}_j \right]$$

*Equation S2*

$$\boldsymbol{s}(x_n) > S(x_{n+1}) \; \forall x_n \, \epsilon \left[\boldsymbol{x}_j \, , \boldsymbol{x}_j + 9\right]$$

*Equation S3*

**Section 3 (Tracking peaks across spectral partitions):** Each spectral partition, $\boldsymbol{S}(:, i)$, is analyzed as follows. The intensity values for each peak region in $\boldsymbol{S}(:, q)$ are extracted and assembled into a matrix $\boldsymbol{\Lambda_q}$, of size 19 × p, where each column, $j$, contains the raw intensity values $\boldsymbol{S}\left(\begin{bmatrix} \boldsymbol{x}_{\boldsymbol{L},j} \\ \vdots \\ \boldsymbol{x}_{\boldsymbol{R},j} \end{bmatrix}, q\right)$ such that

$$\boldsymbol{\Lambda}_q = \left[ \boldsymbol{S}\left( \begin{bmatrix} \boldsymbol{x}_{\boldsymbol{L},1} \\ \vdots \\ \boldsymbol{x}_{\boldsymbol{R},1} \end{bmatrix}, q \right), \boldsymbol{S}\left( \begin{bmatrix} \boldsymbol{x}_{\boldsymbol{L},2} \\ \vdots \\ \boldsymbol{x}_{\boldsymbol{R},2} \end{bmatrix}, q \right), \ldots, \boldsymbol{S}\left( \begin{bmatrix} \boldsymbol{x}_{\boldsymbol{L},p} \\ \vdots \\ \boldsymbol{x}_{\boldsymbol{R},p} \end{bmatrix}, q \right) \right]$$

*Equation S4*

Here, each column in $\boldsymbol{\Lambda}_q$ represents the 19-bin peak region, centered at $\boldsymbol{x}_j$. Similarly, the corresponding bins for the intensity values of the peak regions in $\boldsymbol{S}(:, q)$ are extracted and assembled into a matrix $\boldsymbol{\Lambda}_x, i$, of size 19 × p, such that

$$\boldsymbol{\Lambda}_{x,q} = \begin{bmatrix} \boldsymbol{x}_{\boldsymbol{L},1} & \boldsymbol{x}_{\boldsymbol{L},2} & \cdots & \boldsymbol{x}_{\boldsymbol{L},p} \\ \vdots & \vdots & \ddots & \vdots \\ \boldsymbol{x}_{\boldsymbol{R},1} & \boldsymbol{x}_{\boldsymbol{R},2} & \dots & \boldsymbol{x}_{\boldsymbol{R},p} \end{bmatrix}$$

*Equation S5*

Each peak region $\boldsymbol{\Lambda}_q(:, j)$ is first corrected by subtracting a linear baseline computed using a first-order polynomial fit.

$$B_{i,j}(x) = a + bx, \qquad where \begin{bmatrix} a \\ b \end{bmatrix} = \begin{bmatrix} 1 & \boldsymbol{\Lambda}_{x,i}(1, j) \\ 1 & \boldsymbol{\Lambda}_{x,i}(p, j) \end{bmatrix}^{-1} \begin{bmatrix} \boldsymbol{\Lambda}_i(1, j) \\ \boldsymbol{\Lambda}_i(p, j) \end{bmatrix}$$

*Equation S6*

The baseline is subtracted from the raw peak region, yielding a corrected peak region.

$$\boldsymbol{\Lambda}'_{\boldsymbol{i}}(:, x) = \boldsymbol{\Lambda}_i(:, x) - B_{i,j}(x), \qquad for\ x \in \left[\boldsymbol{\Lambda}_{x,i}(1, j), \boldsymbol{\Lambda}_{x,i}(p, j)\right]$$

*Equation S7*

This baseline-corrected region is then fitted to a Lorentzian function.

$$L(x; x_0, \gamma, A) = \frac{A}{\pi} \frac{05 \times \gamma}{(x - x_0)^2 + (0.5 \times \gamma)^2}, \qquad for\ x \in \left[\boldsymbol{\Lambda}_{x,i}(1, j), \boldsymbol{\Lambda}_{x,i}(p, j)\right]$$

*Equation S8*

where $x_0$ is the peak center, $\gamma$ is the width, and A is the amplitude.

The optimal parameters are determined by minimizing the sum of squared residuals,

$$R(x; x_0, \gamma) = \sum_j [\boldsymbol{\Lambda}'_{\boldsymbol{i}}(:, x) - L(x; x_0, \gamma, A)]^2, \qquad for\ x \in \left[\boldsymbol{\Lambda}_{x,i}(1, j), \boldsymbol{\Lambda}_{x,i}(p, j)\right]$$

*Equation S9*

From each spectral partition, $i$, and peak region, $j$, the fitted centroid, $x_0$, are stored into matrices $\widehat{\boldsymbol{x}}_{\boldsymbol{0}}(j, i)$, of size P×Q, where P is the number of peaks and Q is the number of spectral partitions (see Figure 2A).

**Section 4 (Solving a smooth function of peak position vs. time):** Where $\widehat{\boldsymbol{x}}_{\boldsymbol{0}}$ is the array of fitted centroids from our Lorentzian modeling to the peaks across the spectral partitions, we seek to solve for $\widehat{\boldsymbol{x}}_{\boldsymbol{f}}$, the smooth temporal trend of the centroids across the partitions. To do this, each row of the peak centroid matrix, $\widehat{\boldsymbol{x}}_{\boldsymbol{0}}$, is independently fitted to a smooth function of the mean acquisition time, $\boldsymbol{t}$, (**Figure 2A**, red fitted line). Specifically, for each peak $j$, the following model is used:

$$\widehat{\boldsymbol{x}}_{\boldsymbol{0}}(j,:) \approx \widehat{\boldsymbol{x}}_{\boldsymbol{f}}(j,:) = \sum_{m=1}^{M} \boldsymbol{a}_m \boldsymbol{t}^{b_m}$$

*Equation S10*

Using the $R^2$ calculated from the fit to each row (peak centroid as a function of time), only the rows corresponding to the top 20 $R^2$ values are retained and used in the following section to solve spectral drift as a function of acquisition time. This ensures that the most temporally coherent and well-modeled peaks are used for spectral drift correction.

**Section 5 (Selecting the Basis Set or Reference Spectrum):** Since the peak centroids shift as a function of time, each spectral partition has its own unique set of coefficients used for Equation 1 and Equation 2. To account for the drift, AutoSpect models the coefficients for each partition as a function of acquisition time. This creates three new variables, $\boldsymbol{\rho_1}$, $\boldsymbol{\rho_2}$, and $\boldsymbol{\rho_3}$, of size Q×1, corresponding to the spectral partition specific entries for $P_1$, $P_2$, and $P_3$, respectively, such that:

$$\boldsymbol{\rho_1} = \begin{Bmatrix} \boldsymbol{p_1}_1 \\ \boldsymbol{p_1}_2 \\ \vdots \\ \boldsymbol{p_1}_q \end{Bmatrix}, \quad \boldsymbol{\rho_2} = \begin{Bmatrix} \boldsymbol{p_2}_1 \\ \boldsymbol{p_2}_2 \\ \vdots \\ \boldsymbol{p_2}_q \end{Bmatrix}, \quad \boldsymbol{\rho_3} = \begin{Bmatrix} \boldsymbol{p_3}_1 \\ \boldsymbol{p_3}_2 \\ \vdots \\ \boldsymbol{p_3}_q \end{Bmatrix}$$

*Equation S11*

$$\min_{\rho_1,\rho_2,\rho_3} \sum_{i=1}^{n} \left[\mu_i - \mu_{ref}\right]^2$$

*Equation S12*

**Section 6 (Selecting the Basis Set or Reference Spectrum)**: AutoSpect solves for the coefficients $\boldsymbol{\rho_1}$, $\boldsymbol{\rho_2}$, and $\boldsymbol{\rho_3}$, by first assigning a basis set for comparison. This basis set is a set of peak centroids against which the peak centroids solved as a function of time above can be compared. This set of basis peaks is determined by assigning one of the partitions as a reference. To determine the reference spectrum, total peak region counts per spectral partition are computed and

$$C_j = \sum_{i=1}^{m} \sum_{k=X_{L_i}}^{X_{R_i}} S_{k,j}$$

*Equation S13*

where m is peak number and j is the corresponding spectral partition in the spectral partition matrix S, above.

The spectrum with the fourth-highest total counts is chosen as the reference:

$$S_{ref} = S_{t_{j^*}}, \qquad j^* = \arg max_j C_j , \qquad 4th\ highest\ index$$

*Equation S14*

**Section 7 (Calculating Partition Specific Mass Calibration Parameters):** The optimization problem is solved using nonlinear least squares:

$$\min_{\rho_1,\rho_2,\rho_3} \sum_{i=1}^{n} \left[\mu_i - \mu_{ref}\right]^2$$

*Equation S15*

**Section 8 (Recomputing the Integrated Spectrum):** Peak drift results in blurred features from the integrated spectrum. To correct for this blurring, the integrated spectrum must be recalculated following drift correction. Since each partition is derived from a known number of raw spectra, we let $\mathbf{w} \in \mathbb{R}^{Q\times 1}$ be the vector containing the number of spectra in each partition and define the total number of spectra as $\boldsymbol{W} = \mathbf{1}^{\top}\mathbf{w}$. The drift-corrected integrated spectrum $\hat{\mathbf{s}} \in \mathbb{R}^{N\times 1}$ is then computed as a weighted average of the columns of $\hat{\mathbf{S}}$, with weights proportional to the number of contributing spectra:

$$\hat{\mathbf{s}} = \hat{\mathbf{S}} \cdot \left(\frac{\mathbf{w}}{W}\right)$$

*Equation S16*

The deblurring of spectra following drift correction can be seen in Figure 3 by looking at the bottom row of plots and comparing the original integrated spectrum with the integrated spectrum from drift correction. Although the peaks are narrower for all three regions, the impact, at least in this example, is most noticeable for *m/z* of 18, where it is easily seen that the peak is much more clearly defined.

**Section 9 (Fitting Isolated Peak Regions to obtain the Centroids and Peak Widths)**: Following initial identification, the isolated peak regions are analyzed using Gaussian and Lorentzian fitting approaches. For each peak region, the raw intensity values, $y_i$, and corresponding indices, $x_i$, are extracted along with the corresponding mass-to-charge values for the corresponding ion, $\mu_i^{LUT}$, from the LUT. The region defined by the index corresponding to the point of maximum intensity and a window of ±4 is fitted to a single Gaussian and the $R^2$ calculated.

$$G(x) = Ae^{-\left(\frac{x-x_0}{2\sigma}\right)^2}$$

*Equation S17*

Then only peak regions with $R^2$>0.7 are carried forward for further analyses.

Next, the fitted centroid and width from the Gaussian fitting are used as initial values for fitting the same region to a Lorentzian function. From this, the centroids ($\boldsymbol{x_{fitted}}$) and peak widths ($\boldsymbol{\sigma_{fitted}}$) are saved along with the convergence status and coefficient of determination. To avoid spurious fits, the $\boldsymbol{x_{fitted}}$ is verified to be contained within the windowed region. If it is not, then the peak region and its associated $\boldsymbol{x_{fitted}}$ and $\boldsymbol{\sigma_{fitted}}$ are discarded. Using the $R^2$ from the Lorentzian fitting, all fitted peak regions with $R^2$>0.75 are carried forward. Using Error! Reference source not found.**Equation 2**, the $\boldsymbol{x_{fitted}}$ is converted to $\boldsymbol{\mu_{fitted}}$. Then $\boldsymbol{x_{fitted}}$, $\boldsymbol{\mu_{fitted}}$, and $\boldsymbol{\sigma_{fitted}}$ are stored into column vectors for further refinement.

$$\boldsymbol{x_{fitted}} = \begin{Bmatrix} x_{fitted_1} \\ x_{fitted_2} \\ \vdots \\ x_{fitted_q} \end{Bmatrix}, \qquad \boldsymbol{\mu_{fitted}} = \begin{Bmatrix} \mu_{fitted_1} \\ \mu_{fitted_2} \\ \vdots \\ \mu_{fitted_q} \end{Bmatrix}, \qquad \boldsymbol{\sigma_{fitted}} = \begin{Bmatrix} \sigma_{fitted_1} \\ \sigma_{fitted_2} \\ \vdots \\ \sigma_{fitted_q} \end{Bmatrix},$$

*Equation S18*

**Section 10 (Solving for Peak Broadening as a Function of *m/z*)**: The design matrix $\mathbf{M} \in \mathbb{R}^{n \times 3}$ is defined as:

$$\mathbf{M} = \begin{bmatrix} 1 & \mu_1 & \mu_1^2 \\ 1 & \mu_1 & \mu_2^2 \\ \vdots & \vdots & \vdots \\ 1 & \mu_n & \mu_n^2 \end{bmatrix}$$

*Equation S19*

where each row corresponds to a fitted centroid $\boldsymbol{\mu_{fitted_i}}$, and $n$ is the number of isolated peaks retained.

The polynomial coefficients $\mathbf{c} \in \mathbb{R}^3$ are obtained by solving the following non-negative least squares (NNLS) problem:

$$\boldsymbol{c} = \arg\min_{c \geq 0} \left\| \mathbf{M}\boldsymbol{c} - \boldsymbol{\sigma_{fitted}} \right\|_2^2$$

*Equation S20*

where $\boldsymbol{\sigma_{fitted}} \in \mathbb{R}^3$ is the vector of observed peak widths.

The expected peak width for each centroid $\boldsymbol{\mu}_i$ is then estimated using the fitted polynomial model:

$$\boldsymbol{\sigma_{expected}} = \mathbf{M}\boldsymbol{c}$$

*Equation S21*

To identify outliers with anomalously large peak widths, the relative deviation from the expected curve is computed:

$$i^* = \arg\max_i \left( \frac{\boldsymbol{\sigma}_{\boldsymbol{fitted}_i} - \boldsymbol{\sigma}_{\boldsymbol{expected}_i}}{\boldsymbol{\sigma}_{\boldsymbol{expected}_i}} \right)$$

*Equation S22*

The most extreme outlier $i^*$ is removed, and the polynomial fit is recomputed with the remaining peaks. This process is repeated until only 10 peaks remain (see Figure 4A).

**Section 11 (Iterative Refinement of Mass-to-Charge Axis Recalibration):** Post calibration, the mass-to-charge axis for the detected spectra needs to be recalibrated. The recalibration can be conducted from the integrated spectrum, but the peaks are blurred because the integrated spectrum is the average of drifted peaks. Mass-to-charge axis recalibration is performed after drift correction using the highly resolved peaks.

Using **Equation 2**, the mass axis (*M*) corresponding to the full spectrum is

$$M_i = \{\mu_1, \mu_2, \dots, \mu_n\} = bin2mz\left(\{bin_1, bin_2, \dots, bin_n\};\ P_{1_i}, P_{2_i}, \text{and } P_{3_i}\right)$$

*Equation S23*

where *M* is a vector of size nx1, and *n* is the number of spectral bins in the mass spectrum (e.g., for the icpTOF S2 instrument, n=15616).

Refinement of mass calibration is an iterative process. Each iteration begins with $M_i$ determined by the initial values for the coefficients $P_{1_i}$, $P_{2_i}$, and $P_{3_i}$, and ends with optimized values for the coefficients ($P_{1_{i+1}}$, $P_{2_{i+1}}$, and $P_{3_{i+1}}$) and $M_{i+1}$ using Equation .

At the end of each iteration, the unique values from $M_i$ and $M_{i+1}$ are taken combined

$$M^* = M_i \cup M_{i+1}$$

*Equation S24*

and the combined bin positions using the previous and current parameters are computed using **Error! Reference source not found.**:

$$B_i^* = mz2bin\left(M^*; P_{1_i}, P_{2_i}, \text{and } P_{3_i}\right), \qquad and\ \ B_{i+1}^* = mz2bin\left(M^*; P_{1_{i+1}}, P_{2_{i+1}}, \text{and } P_{3_{i+1}}\right)$$

*Equation S25*

Then, recalibration is considered converged when $B^*_{i+1} = B^*_i$. Figure 4B shows how various regions of the spectra can shift, and that recalibration of the *m/z* axis is not always just a simple translation. As can be seen if Fig. 4B, around both *m/z* of 19 and 203, both regions required a corrective shift to higher *m/z*, while around *m/z* of 60 required a corrective shift to lower *m/z*.

**Section 12 (Dilation of Isolated Peak Regions):** The 10 isolated peak regions, identified in **section** Error! Reference source not found. are adaptively dilated. The left boundary ( $\boldsymbol{x_L}$) of the integration window is initially set at the starting index assigned by the LUT for each identified peak. To adjust this boundary, AutoSpect iteratively examines the mean intensity in a forward window ($\boldsymbol{x_L}$ to $\boldsymbol{x_{L+3}}$) and a backward window ($\boldsymbol{x_{L-3}}$ to $\boldsymbol{x_L}$). If the mean intensity in the forward window exceeds that of the backward window, the boundary is shifted leftward by one unit, i.e., $\boldsymbol{x_L}$à $\boldsymbol{x_{L-1}}$. This process continues until the intensity trend reverses or the boundary approaches the lower limit of the spectrum, at which point further shifting is halted, and the final $\boldsymbol{x_L}$ is assigned.

$$n = \boldsymbol{x_L}(i), \qquad while \;\; \frac{1}{3}\sum_{j=n}^{n+3} S(j) > \frac{1}{3}\sum_{j=n-3}^{n} S(j)\,, \qquad n = n - 1$$

*Equation S26*

Similarly, the right boundary ($\boldsymbol{x_R}$) is initially set at but the LUT and adjusted dynamically. If the mean intensity in the forward window ($\boldsymbol{x_R}$ to $\boldsymbol{x_{R+3}}$) is lower than that in the backward window ($\boldsymbol{x_{R-3}}$ to $\boldsymbol{x_R}$), the boundary is shifted rightward by one unit, i.e., $\boldsymbol{x_R}$ à $\boldsymbol{x_{R+1}}$. This adjustment continues until the trend reverses, or the boundary reaches the upper spectrum limit, at which point further shifting ceases, and $\boldsymbol{x_R}$ is set.

$$n = \boldsymbol{x_R}(i), \qquad while \;\; \frac{1}{3}\sum_{j=n}^{n+3} S(j) < \frac{1}{3}\sum_{j=n-3}^{n} S(j)\,, \qquad n = n + 1$$

*Equation S27*

**Section 13 (Empirical Peak Center Profile via LUT Interpolation):** Given a desired centroid *m/z* denoted as $\mu_0 \in \mathbb{R}^m$, for each of m peaks, the corresponding empirical peak shape is retrieved via interpolation from a precomputed LUT. This process involves several variables, an interpolation operator, and a series of transformation steps:

Variables

$\mu_{0,i}$: the *m/z* centroid for the i[th] peak,

$\overline{b_i}$: the fractional bin index corresponding to $\mu_{0,i}$,

$j_1, j_2$: the nearest lower and upper integer bin indices surrounding $\overline{b_i}$,

$\alpha$: interpolation weight between $j_1$ and $j_2$,

$y_i(s) \in \mathbb{R}^k$: empirical peak profile in standard deviation space,

$x_i(s) \in \mathbb{R}^k$: empirical spectral (non-integer) bin coordinates in standard deviation space,

$b_i(s) \in \mathbb{R}^k$: detector bin numbers in standard deviation space,

$\chi = \{1, \ldots, N\} \in \mathbb{R}$: spectral bin grid

$G \in \mathbb{R}^{N \times m}$: interpolated and normalized peak shape matrix storing interpolated peak shapes for all $m$ peaks.

Interpolation Operator Definition

To interpolate peak shapes from standard deviation space into detector bin space, we define a linear interpolation operator:

$$\mathcal{I}_{x_i \to \chi} : \mathbb{R}^k \to \mathbb{R}^N$$

*Equation S28*

This operator maps a peak shape $y_i(s)$, defined over non-integer spectral bin coordinates $x_i(s)$ in standard deviation space, onto the uniform detector bin grid $\chi$.

**Section 14 (Interpolation Procedure)**

1. Convert *m/z* to fractional bin index using the calibration parameters. Retrieving the empirical peak profile begins by using **Error! Reference source not found.** and g

lobal calibration coefficients ($P_1$, $P_2$, and $P_3$) to calculate the fractional bin index, $\overline{b_\iota}$, corresponding to the theoretical *m/z* for the peak centroid, $\mu_{0,i}$.

$$\bar{b}_i = mz2bin(\mu_{0,i}; P_1, P_2, P_3)$$

*Equation S29*

2. Identify the nearest integer bins and compute weight. Since $\bar{b}_i$ will only extremely rarely be a whole integer, the two nearest corresponding integer bins are determined using flooring and ceiling operators.

$$j_1 = \lfloor \bar{b}_i \rfloor, \qquad j_2 = \lceil \bar{b}_i \rceil, \qquad \alpha = \bar{b}_i - j_1,$$

*Equation S30*

3. Blend adjacent LUT columns to generate intermediate profile corresponding to $\bar{b}_i$. Using the indices $j_1$ and $j_2$, and the fractional weight, $\alpha$, both the intensity profile, $y_i$, and the bin coordinates in standard deviation space, $x_i$, are computed.

$$y_i = (1 - \alpha) \cdot \mathrm{LUT}_y(:, j_1) + \alpha \cdot \mathrm{LUT}_y(:, j_2)$$

*Equation S31*

$$x_i = (1 - \alpha) \cdot \mathrm{LUT}_x(:, j_1) + \alpha \cdot \mathrm{LUT}_x(:, j_2)$$

*Equation S32*

4. Apply interpolation operator to calculate the empirical peak profile. Using $x_i$ and $y_i$, the peak shape $y_i$, defined over non-integer spectral bin coordinates $x_i$, in standard deviation space, is mapped onto the uniform detector bin grid $\chi$.

$$G(:, i) = \mathcal{I}_{x_i \to \chi}\,(y_i)$$

*Equation S33*

5. Normalize the empirical peak profile to unit area.

$$G(:, i) \leftarrow \frac{G(:, i)}{\sum G(:, i)}$$

*Equation S34*

This yields $G \in \mathbb{R}^{N \times m}$, the empirical core peak shapes for each centroid $\mu_{0,i}$.

**Section 15 (Tail Widths Constrained by Peak Width):** Tailing widths, $\boldsymbol{\beta}$, follow a smooth dependence on peak resolution across the mass range. As such, tail widths are not free parameters; rather, they are scaled by the peak widths according to a first-order polynomial in centroid *m/z*.

Let:

- $\mu \in \mathbb{R}^m$: centroid *m/z* values,
- $\mathbf{M} \in \mathbb{R}^{m\times 3}$: design matrix with rows [1, $\mu_0$],
- $\mathbf{c} \in \mathbb{R}^3$: polynomial coefficients.
- $\mathbf{R} \in \mathbb{R}^{\boldsymbol{m}}$: vector of resolution-scaling factors,
- $\boldsymbol{\beta} \in \mathbb{R}^{\boldsymbol{m}}$: vector of asymmetric tailing widths.

Then:

$$\mathbf{R} = \mathbf{M} \cdot \boldsymbol{c}$$

*Equation S35*

$$\boldsymbol{\beta} = \mathbf{R} \circ \boldsymbol{\sigma}$$

*Equation S36*

**Section 16 (Mass-to-Charge Dependent Tail Amplitude Scaling):** The amplitude of the low- and high-*m/z* tails are on the order of 0.0001 the intensity of the parent peak and vary smoothly across the spectrum. We have modeled this dependence as an exponential.

Let:

- $\mu_{0,i} \in \mathbb{R}$ represents the centroid position in *m/z* units;
- $R = [R_1, R_2, R_3] \in \mathbb{R}^3$ are empirical coefficients.

Then, for each peak centroid position $\mu_{0,i}$, the tail amplitude ratio $\boldsymbol{r}_i$ is computed using:

$$\boldsymbol{r}_i = R_3 \cdot \exp(R_2 \cdot \mu) - R_1$$

*Equation S37*

This scaling function is applied independently to the low- and high-energy tails to produce the vectors $\boldsymbol{r}_{\mathbf{low}}$ and $\boldsymbol{r}_{\mathbf{high}}$, which are then used below during peak shape assembly. This approach allows the contribution of asymmetric tails to increase or decrease systematically with mass, improving overall peak modeling fidelity across a broad dynamic range.

**Section 17 (Peak Shape Combination and Normalization):** After empirical peak shapes and asymmetric tails have been defined, they are combined into a final asymmetric model for each elemental signal. This step ensures that each modeled peak incorporates core instrument resolution and realistic distortion due to ion scattering and detector overload.

Let:

- $\mathrm{G} \in \mathbb{R}^{N\times m}$: empirically determined profiles of the peak centers (interpolated from $LUT_x$ and $LUT_y$), where $m$ is the number of total spectral components
- $\mathbf{T_{low}}, \mathbf{T_{high}} \in \mathbb{R}^{N\times m}$: low- and high- *m/z* tailing components (scaled by peak width σ),
- $\mathbf{r_{low}}, \mathbf{r}_{\boldsymbol{high}} \in \mathbb{R}^{1\times m}$: amplitude scaling ratios for the low-μ and high-μ tails.

Because both tail matrices are normalized, their total area is 1. Tails are then rescaled by the ratio of the empirical peak height to the maximum tail intensity and then scaled by the desired amplitude ratio. The asymmetric peak model is:

$$\mathrm{G_T} = \mathrm{G} + \mathrm{T_{high}} \circ \left(\frac{\boldsymbol{r}_{\mathbf{high}} \circ \max(\mathrm{G})}{\max\left(\mathbf{T_{high}}\right)}\right) + \mathrm{T_{low}} \circ \left(\frac{\boldsymbol{r}_{\mathbf{low}} \circ \max(\mathrm{G})}{\max(\mathbf{T_{low}})}\right)$$

*Equation S38*

After combining the core peak with the asymmetric tails, each resulting peak shape $G_T(:,i)$ is normalized such that its total area integrates to one (Equation ).

The resulting matrix $\mathrm{G_T} \in \mathbb{R}^{N\times m}$ contains the asymmetric, resolution-constrained, and normalized peak shapes for each elemental signal prior to application of instrumental notching and isotopic branching.

**Section 18 (Correction for Instrumental Notching, Tofwerk-Specific):** AutoSpect includes an optional correction mechanism specific to Tofwerk TOF-MS instruments, which employ inverse quadrupole filtering, referred to as notching, to reduce the transmission of extremely abundant background ions, such as those arising from argon, nitrogen, water vapor, and other atmospheric components, which would otherwise cause serious detector saturation. This filtering alters the transmission efficiency at selected *m/z* values, artificially suppressing the observed intensities of certain isotopes and thereby distorting their natural abundance ratios.

To correct for this effect, AutoSpect uses the stored notching parameters to automatically optimize and account for these changes. These values are used to compute a set of scaling factors, referred to as FLOAT, which are applied to the branching ratios of affected isotopes.

Let:

- $\mathbf{B}_{\text{iso}} \in \mathbb{R}^{m\times\gamma}$: original branching ratio matrix before notching correction,
- $\mathbf{F} \in \mathbb{R}^{\gamma\times\gamma}$: diagonal matrix of scaling factors (with ones for unaffected isotopes),
- $\widetilde{\mathbf{B}}_{\text{iso}} = \mathbf{B}_{\text{iso}} \cdot \mathbf{F}$: corrected branching ratio matrix used in downstream modeling.

The $\mathbf{F}$ matrix adjusts only a small number of isotopes (typically four), and notching parameters are stored alongside the dataset for full reproducibility.

**Section19 (Spectral Fitting and Peak Deconvolution):** In the final steps of AutoSpect, using the drift calibration and the branched equations defined in the previous sections, the spectra for the samples, gas blanks, and reference standards are fitted to deconvolute peak overlap.

Let:

- $\mathbf{x}_q \in \mathbb{R}^{N}$: drift-corrected mass axis for partition $q$
- $\mathbf{F}_q \in \mathbb{R}^{N\times M_q}$: matrix of observed spectra for partition $q$
- $\mathbf{B} \in \mathbb{R}^{N}$: background model defined on the reference axis
- $\mathbf{B}_q \in \mathbb{R}^{N\times M_q}$: interpolated background per spectrum for partition $q$
- $\mathbf{C}_q \in \mathbb{R}^{q\times M_q}$: output matrix of fitted coefficients for partition $q$
- $\tilde{G}_q \in \mathbb{R}^{N\times\gamma}$: interpolated modeled isotope-resolved peak shapes for partition $q$

Interpolation Operator Definition

Let:

- $\mathbf{x}_0 = \left[x_{\mathbf{0}}^{(\mathbf{1})}, x_{\mathbf{0}}^{(\mathbf{2})}, \ldots, x_{\mathbf{0}}^{(N)}\right]^{\top} \in \mathbb{R}^{\boldsymbol{N}}$: the reference mass axis
- $\mathbf{x}_q = \left[x_q^{(\mathbf{1})}, x_q^{(\mathbf{2})}, \ldots, x_q^{(N)}\right]^{\top} \in \mathbb{R}^{\boldsymbol{N}}$: the drift-corrected axis for partition $q$
- $M \in \mathbb{R}^{\boldsymbol{N} \times \boldsymbol{q}}$: a matrix whose columns $M_j$ are defined on $\mathbf{x}_0$

We define the interpolation operator $\mathcal{I}_{x_0 \to x_i}(M)$ as:

$$\left[\mathcal{I}_{x_0 \to x_q}(M)\right]_j = P_j(\mathbf{x}_q)$$

*Equation S39 1*

where $P_j(x)$ is the piecewise cubic Hermite interpolating polynomial fitted to the column vector $M_j$ on the support $\mathbf{x}_0$computed using the Fritsch–Carlson method.

Step 1: Interpolate the Model Matrix

Typically, with LA data, the individual pixel spectra are sparsely populated where most of the spectral bins contain zeros. This is especially the case with biological tissues. Because of this, interpolating the pixel spectra into a common grid for drift correction can result in significant errors. To avoid such potential errors, the composite model matrix $G_{br}$ is interpolated to the local drift-corrected mass axis for each partition:

$$\tilde{G}_q = \mathcal{I}_{x_0 \to x_q}(G_{br})$$

*Equation S402*

Step 2: Background Subtraction

Prior to spectral fitting, background correction is applied using either measured gas blanks or an automated SNIP-based background estimation algorithm.

(a) Gas Blank Subtraction

When dedicated gas blank spectra are available, the gas blank spectra are averaged following drift correction and then interpolated into the *m/z* frame of the partition being corrected. As such, the background is interpolated from the reference axis to the drift-corrected mass axis:

$$\mathbf{B}_i{}^{(k)} = \mathcal{I}_{x_0 \to x_i}(\mathbf{B})$$

*Equation S413*

(b) SNIP Algorithm [22]

In the absence of gas blanks, AutoSpect employs the **S**tatistics-sensitive **N**on-linear **I**terative **P**eak-clipping (SNIP) algorithm, adapted from Ryan et al. (1988) to estimate background. This procedure performs robust baseline estimation via the following steps:

1. Logarithmic Transformation:

The input spectrum $y$ is smoothed using a nested logarithmic transformation to reduce peak dominance:

$$y' = \log(\log(y + 1) + 1)$$

*Equation S424*

2. Theoretical Peak Width Estimation:

A dynamic window width $w_i$ is computed for each spectral channel, $i$, using the expected standard deviation $\sigma_{expected_i}$ of the peak shape (as modeled in **Section 4.4**). This accounts for the natural mass-dependent broadening of peaks.

3. Iterative Peak-clipping:

   The transformed spectrum is subjected to a number of iterative clipping passes. At each iteration $j$, a smoothed background estimate is computed using symmetric windows of width $w_i$, and values above the local average are clipped:

$$y_i^{(j+1)} = \min\left( y_i^{(j)}, \frac{y_{i-w_i}^{(j)} + y_{i+w_i}^{(j)}}{2} \right)$$

*Equation S43 5*

4. Inverse Transformation:

   After the final iteration, the background estimate is reconstructed using the inverse of the nested logarithm:

$$B_q{}^{(k)} = \exp\left(\exp\left(y_i^{(final)}\right) - 1\right) - 1$$

*Equation S44 6*

Step 3: Solve Linear System

Once the model matrix and background are aligned with the measured data, the corrected spectra are deconvolved using standard linear least squares:

$$\mathbf{C}_q(:,k) = \left(\tilde{G}_q^\top \tilde{G}_q\right)^{-1} \tilde{G}_q^\top \left(\mathbf{F}_q(:,k) - \mathbf{B}_q(:,k)\right)$$

*Equation S45 7*

This yields $\mathbf{C}_q$, a matrix of amplitudes for all components (columns of $\tilde{G}_q$) across all spectra in partition $q$. Since each column of $\tilde{G}_q$ is normalized to unit area, the resulting coefficients in $\mathbf{C}_q$ represent the total extracted counts for each modeled peak component, and no additional normalization or post-processing is required.

**Section 20 (Derivation of TOF-MS Calibration Function)**: In a time-of-flight mass spectrometer (TOF-MS), ions are accelerated through an electric potential and then travel through a field-free drift region before striking the detector. The time t that an ion takes to reach the detector is determined by its mass-to-charge ratio (m/z), and is derived from the classical kinetic energy equation:

$$K_E = \frac{1}{2}mv^2 = qV$$

*Equation S46*

where m is the ion mass, v is the velocity, q is the ion charge, and V is the accelerating voltage. Solving for v and expressing the time-of-flight as t = L/v, where L is the flight path length, gives:

$$t = L\sqrt{\frac{m}{2qV}} = L\sqrt{\frac{m/z}{2V}}$$

*Equation S47 8*

This yields a square-root dependence of t on m/z, consistent with the idealized flight model. However, to account for instrumental delays and nonlinearities in practical TOF-MS systems, this relationship is generalized as:

$$t = P_1 \left(\frac{m}{z}\right)^{P_3} - P_2$$

*Equation S48 9*

where $P_1$, $P_2$, and $P_3$ are empirical calibration constants determined from known reference peaks. This formulation is used in AutoSpect to interconvert between digital time bins and the corresponding calibrated mass-to-charge values.
